# Optimization of Microwave-Assisted Extraction of Antioxidant Compounds from *Vitex negundo* Leaves using Response Surface Methodology

**DOI:** 10.1101/2025.03.15.643432

**Authors:** Anindita Kundu, Niladri Sett

## Abstract

**Introduction:** Polyphenolic substances are obtained from the leaves of *Vitex negundo* has gained attention for its therapeutic benefits.

**Objectives:** To obtain the maximized antioxidant-rich polyphenolic substances from the *Vitex negundo* leaves via microwave-assisted extraction (MAE).

**Material and Methods:** Box–Behnken design (BBD) was employed to evaluate the influence of several variables of MAE on the total phenolic content (TPC) and total flavonoid content (TFC) as determined by Folin–Ciocalteu assay method and Aluminum Chloride assay method, and the antioxidant capacity as measured by DPPH and ABTS assay methods, respectively. Response surface methodology (RSM) was used for finding optimal extraction conditions. GC-MS analysis was conducted on the extract. The tocopherol content and anticancer potential were also estimated using HPLC and MTT assay respectively.

**Results:** The ideal extraction parameters were found to be 14 minutes, 48 °C, 65 % (v/v) methanol concentration, and 14 mL of extraction solvent. The optimum experiment conditions produced the TPC and TFC values of 1.46 mg GAE/g extract and 1.06 mg QE/g extract respectively. Furthermore, the DPPH and ABTS assays results showed the optimum values of 53 % and 77 % respectively. 12 bioactive compounds were identified using GC-MS. The amount of tocopherol was found to be 414.87 µg/g. Lastly, the obtained leaf extract was demonstrated its anticancer potential on PC3 cell lines.

**Conclusion:** The findings demonstrated leaf extract’s potential as a useful source of polyphenols with strong antioxidant qualities that can be used in a variety of pharmaceutical applications.

## 1 Introduction

Since prehistoric times, herbal medicines have been receiving great attention as potential therapeutic agents for the management of human health care and well-being [1]. Research evidence reveals that 69% of new antibacterial agents (1981–2006) and 49% of new anticancer agents (1940–2014) in marketed products are derived from natural products. This highlights the significant role of natural sources in drug discovery [2]. In the present circumstances, the employment of herbal medicinal principles is getting worldwide acceptance with a rapid dissemination of innovative technologies because of having fewer unwanted effects, low price and potential to overcome the limitations of conventional medicine [1].

*Vitex negundo* is the one of the important aromatic medicinal plants, and is widely used as folk medicine for the management of several health problem such as microbial attack, inflammation, CNS-depression, tumour, cardiac issues, oxidative stress, fluid retention in body, cancer and liver disorders, etc [3], [4]. *Vitex negundo* is a deciduous aromatic shrub and distributed worldwide including Afghanistan, Pakistan, India, Sri Lanka, Bangladesh, Bhutan,China, Indonesia, Japan, Korea, Kenya, Madagascar, Malaysia, Mozambique, Myanmar, Nepal, Taiwan, Tanzania, Thailand and Vietnam [5]. It has been reported that the essential oil of *Vitex negundo* leaves chiefly consists of monoterpenes, sesquiterpenes, various fatty acids etc[6] [7].

In current research trends, polyphenols and vitamin E are extensively studied for the promising antioxidant properties, and also used in adjuvant therapy in carcinoma [8]. However, ATBC trials revealed that the α-tocopherol alone and together with β-carotene could play a key role to reduce the death risk in prostate cancer[9]. Awad *et al*. [10] explored the importance of dietary phytosterols especially β-sitosterol in prevention of growth and dissemination of prostate cancer cells [11]. It is already known that the terpenoids, the monoterpenes and sesquiterpenes are the major constituents of essential oil [12]. Research findings suggested the significant employment of dietary terpenoids as chemo-preventive agents at pre-malignant stage and various developmental stages of cancers; and especially in prostate carcinoma by down-regulating the Bcl-2 protein as well [13]. Existing literatures demonstrated that the terpenes and their derivatives such as citral, squalene, thunbergol, phytol and phytosterol such as β-sitosterol, γ-sitosterol are the major phytoconstituents of essential oil; and they are considered to be responsible for possessing strong antioxidant property[14], [15].

An earlier study [14], the methanol extract and the essential oil of *Vitex negundo* leaves had been evaluated for β-sitosterol and polyphenol contents using HPLC screening method; finally, the β-sitosterol and thirteen different polyphenols were identified with different quantities. The leafs extracts were also showed the significant anti-inflammatory potentials on the RAW 264.7 cells [16].

Several research have optimized the extraction parameters to get the maximum target bioactive compounds from the extract of the *Vitex negundo* leaves. For those, the extraction conventional methods were used including the soxhlet extraction, hot solvent extraction and ultrasonic extraction. Mostly, the various solvents (ethanol-aqueous, methanol, ethanol) were used with different times and temperature for optimization [17], [18], [19]. Nowadays, ultrasonic extraction method has become more well known because it may enhance extraction yields, decrease solvent usage, and minimize extraction times [20], [21]. Ultrasonic extraction works by using cavitation, vibration, crushing and mixing produced by the ultrasonic waves, that increase the mass transfer of the bioactive materials by rupturing the plant cell walls [22]. But this system has some drawbacks including high cost and unintended alterations of the molecules.

Microwave-assisted extraction (MAE) is an ecologically friendly extraction method that utilizes microwaves to extract materials that have been soaked in a solvent [23], [24]. Unlike traditional extraction techniques, microwave radiation may heat the reactants and solvent directly by penetrating the wall of the extraction vessel [25]. Because of having several benefits, MAE is extensively employed in the laboratory. It requires low energy, less amount of solvent and reduces the waste production. It has drawn a lot of interest in both scientific study and industrial applications because to its effective extraction of bioactive compounds, which makes it a useful tool in the field of natural product extraction [26], [27]. The intricacy of mass transfer and restricted penetration depth of microwave irradiation can be affected by some parameters like temperature, microwave frequency, current obstacle to scale up the MAE process. Thus, scaling up MAE for commercial uses needs a thorough examination in which different factors influence extraction kinetics. Therefore, it is essential to develop models which can estimate the ideal experimental conditions [28], [29].

In this research, we evaluated the phytoconstituents and the antioxidant potentials of *Vitex negundo* leaves, targeting to uncover the benefits in pharmacological uses. To achieve this goal, we used optimization technique based on MAE for the identification and quantification of the bioactive compounds from the *Vitex negundo* leaves. The response surface methodology (RSM) was adopted using a Box–Behnken design (BBD) as the experimental design, and the model fit, regression equations, analysis of variance and three-dimensional response surface plots were generated. Temperature (50-80 °C), extraction time (5-25 min), extraction volume (5-15 mL) and methanol concentration (40-80 % v/v) were examined as the primary factors influencing the efficiency of extraction and the antioxidant activities. Total phenolic content (TPC), total flavonoid content (TFC), DPPH concentration, ABTS contration results were recorded by implementing the Box–Behnken design. GC-MS platform was used for identifying the bioactive compounds from the leaf extract which was produced under the ideal extraction settings identified by the proposed model. HPLC technique was employed to estimate the tocopherol content presented in the optimal leaf extract. Finally, *in-vitro* anticancer property of optimal leaf extract was assessed on a prostate carcinoma cell line.

## 2 Experimental

### 2.1 Materials

Methanol was purchased from Spectrochem Pvt. Ltd., Mumbai, India; gallic acid, quercetin, DPPH, ABTS, tocopherol from HiMedia Laboratories Pvt. Ltd., Mumbai, India, and Sigma-Aldrich. Folin-denis reagent was purchased from Sigma Aldrich Co., St. Louis, MO, USA. Dulbecco’s Modified Eagle’s Medium (DMEM) was procured from Gibco, Grand Island, NY 14072, USA1-716-774-6700. All the chemicals and reagents used in this research were of high analytical grade and obtained from Hi-Media Research Laboratories Pvt. Ltd., Mumbai, India.

### 2.2 Preparation of plant extract for microwave assisted extraction (MAE)

Fresh leaves of *Vitex negundo* were collected from the nursery of the Maulana Azad College (Kolkata; West-Bengal the state of India). Then, the leaves were shade dried for 15 days and subjected for size reduction by a cutting mill. To acquire a uniform plant matrix, the small sections of leaves were passed through a 60-mesh size sieve and then kept in ziplock pouches for the experiment.

The extraction system comprised of microwave extractor (CATA R) manufactured by Catalyst Systems (Pune, India) equipped with a magnetron of 2450 MHz with a maximum power of 700 W (100 %), a reflux unit, 10 power levels (140 W (20 %) to 700 W (100 %)), time controller, temperature sensor, exhaust system, beam reflector and a stirring device.

The experiments were performed using a microwave extractor coupled with a high efficiency condensing unit. 1 g of dried and finely powdered leaf was precisely weighed and imbibed to methanol for 15 min. The extraction was conducted with an intermittent way of irradiation for 1 min, followed by cooling for 1 min. The inbuilt condensing unit makes the extraction more effective and robust. After extraction, the extract was passed through No. 1 Whatman filter paper and then the solvent was evaporated by rotary vacuum evaporator which was centrifuged at 82 rpm, 43 °C. Afterwards, the dried residue was collected and reconstituted with methanol and stored for further qualitative and quantitative analysis [30].

### 2.3 Response surface methodology modelling

Response surface methodology was employed to assess the optimal conditions for the microwave assisted extraction to get the highest amounts of total phenolic content, total flavonoid content and antioxidant potentials (DPPH free radical scavenging capacity, ABTS radical scavenging capacity) of leaf extract of *Vitex negundo*. The correlation between four independent variables (A: temperature, 50–80 °C; B: time, 5–25 min; C: extraction volume, 5– 15 mL; D: methanol concentration, 40––80 % v/v) and the response variables of total phenolic content (TPC, *Y1*), total flavonoid content (TFC, *Y2*) and scavenging potentials (DPPH free radical scavenging assay, *Y3* and ABTS radical scavenging assay, *Y4*) was determined applying box-behnken design. All independent variables were linked to the different coded variable levels (-1, 0, +1) (Table 1). There were 27 experimental runs created, comprising three central points. All the experiments were carried out at random, and the range of studied variables was chosen based on preliminary testing and experimental constraints. Each analysis was carried out in three times to establish the reliability and reproducibility. Results were expressed as mean ± standard deviation (SD). RSM was conducted using the “rsm”^1^ library of the statistical programming language “R”^2^. The experimental results obtained, were used to fit quadratic polynomial regression models based on the following equation:

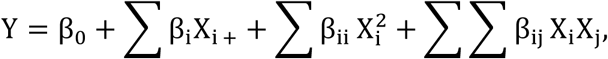

**Table 1.** Selected extraction parameters for BBD optimization.

| Independent variable | Symbol | Factor level |  |  | Dependent variable |
| --- | --- | --- | --- | --- | --- |
|  |  | -1 | 0 | +1 |  |
| Temperature (°C) | A | 50 | 65 | 80 | Y1: TPC (mg GAE/g extract)<br>Y2: TFC (mg QE/g extract)<br>Y3: %DPPH<br>Y4: %ABTS |
| Time (min) | B | 5 | 15 | 25 |  |
| Extraction Volume (mL) | C | 5 | 10 | 15 |  |
| MeOH (%) | D | 40 | 60 | 80 |  |

where *Y* is the response variable, each of *X_i_* and *X_j_* is an independent variable, *β*_0_, *β_i_, β_ii_* and *β_ij_* are the intercept coefficient, the linear coefficients, the quadratic coefficients, and the interaction coefficients of *X_i_* and *X_j_* respectively.

The response surface and contour plot approaches were employed to demonstrate the connection between responses and various degrees of independent variables and forms of interaction between two independent variables. The ultimate confirmation experiment (*n* = 3) was conducted utilizing optimized independent extraction variables, and the experimental results that were compared with predicted values for validation of the model. Analysis of variance (ANOVA) test was performed, and the maximum *R^2^* and adjusted *R^2^* values were employed to evaluate the accuracy of the estimated coefficients, and *p-*values ≤ 0.05 were considered significant. Optimum experimental conditions for MAE were obtained by simultaneously optimizing the four response variables using desirability function approach (Derringer & Suich, 1980).

### 2.4 Determination of total phenolic content (TPC)

A preliminary test for total phenolic content of leaf extract of *Vitex negundo* plant was executed using the Folin–Ciocalteau method as proposed by Chouhan et al., with a little modification [17]. Briefly, aliquot of sample (mg/mL) was added in 4 mL of sodium carbonate solution (75 g/L) and 5 mL of 10 % Folin-Ciocalteu reagent. Equation of the calibration curve was prepared (y = 0.136x + 0.001, R^2^ = 0.999) by using the gallic acid (with a range of 10 to 80 μg/mL) as the standard, and whereas the methanol was used as a blank. Then the prepared mixture was vortexed and kept in incubation for 1 hour in dark place and immediately the absorbance was measured at 765 nm using a UV-Vis spectrophotometer. The prepared mixture was measured for the three replicates of the sample. Finally, the results were expressed as gallic acid equivalents in milligram per gram (mg GAE/g) of dried extract.

### 2.5 Determination of total flavonoid content (TFC)

The total flavonoid content was quantitatively determined using the aluminium chloride colorimetric assay method with slight modifications as narrated by Chouhan et al. [31]. In brief, leaf extract (mg/mL) was diluted with 300 μL 5 % NaNO_2_ solution, 300 μL of 10 % AlCl_3_ solution, 4 mL distilled water, and was allowed to incubate for 6 min. Thereafter, 2 mL of 1 M NaOH solution was added to the mixture and allowed to stand still the solution for 15 min and then checked the absorbance at 510 nm using a UV-Vis spectrophotometer. The total flavonoid content was calculated on the basis of calibration curve (y = 0.097x + 1.489, R^2^ = 0.994) obtained by using quercetin (ranging from 10 to 80 μg/mL) as the standard solution and whereas the methanol was used as a blank. The prepared mixture was measured for the three replicates of the sample. The result was expressed as equivalents in milligram per gram (mg QE/g) of dried extract.

### 2.6 Antioxidant assays

#### 2.6.1 DPPH free radical scavenging assay

The free radical scavenging activity of leaf extract was assessed by the DPPH free radical scavenging method with little modifications to the conditions previously described by Chouhan et al. [31]. 0.2 mL of the sample (100 μg/mL) was mixed with 2 mL of 0.1 mM 2,2-diphenyl-1-picrylhydrazyl (DPPH) stock solution in methanol, and left to incubate for 30 min in a dark place. Ascorbic acid was used as positive control and prepared in the same procedure. After the incubation period the absorbance was recorded against the methanol as a blank solution at 517 nm using a UV-Vis spectrophotometer. The percentage of inhibition of DPPH was reported in comparison with the control solution (containing all of the reagents and methanol instead of the test sample). This assay was carried out in triplicate to establish the reliability and the inhibitory potential DPPH was calculated as the following equation:

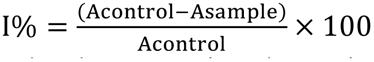

In the above-mentioned equation, A_control_ is the absorbance of the control solution, and A_sample_ is the absorbance of a sample solution.

#### 2.6.2 ABTS radical scavenging assay

The ABTS radical scavenging activity assay was executed with little modifications of method which was reported by Ondua et al. [32]. 7.4 mM of ABTS was mixed with 2.6 mM of potassium persulphate in an equal proportion. Then the prepared stock solution was kept in a dark place at room temperature for 12-16h. Thereafter, 1 ml. of ABTS stock solution was mixed with different concentrations (20-100 μg/ml) of 1 ml of sample solutions (methanol extract and essential oil) and incubated in a dark place for 2 hours. Finally, the absorbances of reaction mixtures were checked at 734 nm with the UV-Vis spectrophotometer. Ascorbic acid was used as positive control and produced in the same fashion. Potassium persulphate was used as the blank solution. This assay was carried out in triplicate to establish the reliability and the inhibitory potential of ABTS was determined using the equation is given below:

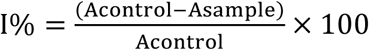

In the aforementioned equation, A_control_ is the absorbance of the control solution, and A_sample_ is the absorbance of a sample solution.

### 2.7 GC-MS conditions

The components of leaf extracts were analyzed by GC equipped with a mass spectrometer detector and performed with a few modifications of method as earlier described by Tohidi et al. [12]. GC-MS analysis was carried out using an Agilent 19091S-433UI gas chromatograph coupled with a HP-5ms 5 % phenylmethylsiloxane capillary column (30 m × 0.25 mm i.d.; film thickness 0.25 μm). Aliquot of 1 μL was injected into the GC injector port with the split ratio 1:40. Initially, the oven temperature was set at 60 °C for 5 mins. Afterwards, the temperature was increased to 240 °C with a successive increment rate of 3 °C/min and followed by at 270 °C with a ramp of 5 °C/min and finally kept constant for 8.5 min. Helium gas was considered as carrier gas and maintained a constant flow rate of 1 mL/min with the ionization voltage of 70 eV. The volatile organic components (VOC) were determined by matching their retention indices (RI, HP-5) with the database of NBS75K library. The name, structure and molecular weight of the identified VOC were comprehended with conductance of the National Institute of Standard and Technology (NIST) covering an extensive range of patterns stored in the library.

### 2.8 Estimation of tocopherol

The estimation of tocopherol of leaf extract of *Vitex negundo* was evaluated by High-performance liquid chromatography (Alliance model 2695) with a UV-Vis detector (2487). Tocopherol was separated by using C18 column (4.6 mm × 150 mm × 5μm particle size), maintained at 30 °C. Tocopherol was eluted out using a mixture of mobile phase, i.e. methanol and water (98:2 v/v) at a constant flow rate of 0.5 mL/min and total run time 15 min. 10 μL of sample (100 μg/mL) was injected into the injector port and detection was carried out at 292 nm. The compound was detected and quantified compared with the peak area and time of the authentic standard (tocopherol) purchased from the Sigma Aldrich dissolved in methanol and diluted in the same manner [33].

### 2.9 *In-vitro* anticancer assay

#### 2.9.1 Cell lines and cell culture conditions

Prostate cancer cell line (PC-3) and normal lung tissue cells (WI-38) were procured from American Type Culture Collection (ATCC) and cultured in the laboratory in DMEM supplemented with 10 % FBS (Invitrogen), 1 % L-Glutamine, 0.1 mM non-essential amino acid, and 100 U/ml penicillin/streptomycin at 37 °C; in a humidified atmosphere with 5 % of CO_2_ [34]. The culture medium was replaced with the fresh medium at every 48 to 72 h. At the time cells reached density, the culture medium was removed from the 75 cm^2^ flask and washed with phosphate-buffered saline (PBS). Afterwards, for removal of adhesion-proteins, the 1.5 mL of Trypsin-EDTA was added and placed into the incubator at 37 °C in a humidified atmosphere with 5 % of CO_2_ for 5 mins. Subsequently, 3 mL of culture media was given to terminate the action of trypsin. Then cells were transferred to a new flask for their sub-culture growth. Finally, cells were quantified by using a haemocytometer.

#### 2.9.2 Cell viability assay

The effect of plant extract on cellular proliferation was evaluated with cell viability assay using the tetrazolium salt (3-[4,5-dimethylthiazol-2-yl]-2,5diphenyl tetrazolium bromide) (MTT) which was reduced by the mitochondrial succinate dehydrogenase and form the formazan crystal [34]. Cultured cells were grown until they reached 70 % of confluence and then treated with the several concentrations (6.25 μg/mL, 12.5 μg/mL, 25 μg/mL, 50 μg/mL, and 75 μg/mL) of leaf extracts or DMSO as vehicle in triplicate. After 24 h and 48h of incubation period the cell viability was assessed by adding MTT solution (5 mg/mL in PBS buffer) into each well and incubated at 37 ^◦^C for 3 h. Then, the cell culture media was removed from each well and newly formed formazan complexes were solublized in 200 µL DMSO. Finally, the absorbance was checked at 595 nm using micro-plate reader and values of cell viability in different concentrations of samples were quantified and plotted on a graph. Different time points (24 h and 48 h) were evaluated during the experiment.

### 2.10 Statistical analysis

All the experiments were executed in triplicate. The results were expressed as means ± SD (*n* = 3) and the statistical analyses were performed using one-way analysis of variance (ANOVA), followed by Bonferroni’s multiple comparison test was performed using GraphPad Prism (version 5.04) software. Statistical differences were considered to be significant at \**p* < 0.05, \*\**p* < 0.01, \*\*\**p* < 0.001 and \*\*\*\**p* < 0.0001.

## 3 Results and Discussion

In this research, we aimed to investigate a contributory role of *Vitex negundo* plant in the health-care system, particularly focusing on the leaves collected in the spring time. The goal of this study was to explore the potential of the leaf of *Vitex negundo* as the natural source of antioxidants.

To get the maximum antioxidant potentials, the extraction parameters of MAE were optimized using Response Surface Methodology (RSM) with the Box–Behnken Design (BBD) as the experimental design with a run of 27. This research considered the impact of four different extraction parameters including temperature, time, extraction volume and methanol concentration. Table 2 demonstrates the experimental design for BBD with experimental and predicted values of TPC, TFC, DPPH and ABTS. Here, the range of TPC values are from 1.08 to 1.42 mg GAE/g extract whereas the range for TFC values are from 0.78 to 1.03 mg QE/g extract. For %DPPH, the range is from 37.18 to 52.75 while the values for %ABTS is from 63.96 to 77.56. The experiments corresponded with three central points (runs: 6, 14, and 23), having mean values of 1.39 ± 0.13 mg GAE/g extract for TPC, 1.00 ± 0.02 mg QE/g extract for TFC, 50.87 ± 0.85 for %DPPH and 76.49 ± 0.54 for %ABTS, respectively. The quadratic equations obtained from the multiple regression analyses, depicting the effect of each response variable against the independent variables: A, B, C, and D corresponding to temperature, extraction time, extraction volume, and methanol concentration respectively, are as follows.

TPC = 1.39 - 0.07 × A - 0.05 × B + 0.05 × C - 0.12 × A^2^ - 0.08 × B^2^ - 0.05 × AC - 0.08 AD + 0.05 × BC + 0.04 × BD
TFC = 1 - 0.05 × A - 0.02 × B + 0.04 × C - 0.1 × A^2^ - 0.05 × B^2^ - 0.05 × AC - 0.06 × AD + 0.05 × BC + 0.03 × BD
DPPH = 50.87 – 3.29 × A - 2.43 × B + 2.81 × C - 6.02 × A^2^ - 2.31 × B^2^ - 2.73 × AC - 3.8 × AD + 3.27 × BC + 1.96 × BD
ABTS = 76.49 - 2.8 × A -1.61 × B + 1.15 × C - 4.04 × A^2^ - 2.18 × B^2^ -1.64 × C^2^ - 1.55 × AB - 2.26 × AC - 2.25 × AD

**Table 2:** Extraction parameters for BBD with experimental and predicted values of TPC, TFC, DPPH and ABTS.

|  | Factors |  |  |  | Y1: TPC (mg GAE/g extract) |  | Y2: TFC (mg QE/g extract) |  | Y3: %DPPH |  | Y4: %ABTS |  |
| --- | --- | --- | --- | --- | --- | --- | --- | --- | --- | --- | --- | --- |
| Run | A | B | C | D | Predicted | Experimental | Predicted | Experimental | Predicted | Experimental | Predicted | Experimental |
| 1 | 0 | 0 | -1 | 1 | 1.28 | 1.29 | 0.92 | 0.92 | 46.89 | 47.23 | 74.17 | 74.50 |
| 2 | 1 | -1 | 0 | 0 | 1.19 | 1.20 | 0.84 | 0.86 | 42.28 | 42.19 | 70.64 | 71.56 |
| 3 | 0 | 1 | 0 | -1 | 1.21 | 1.21 | 0.88 | 0.86 | 42.79 | 42.95 | 71.85 | 72.01 |
| 4 | -1 | 0 | 0 | -1 | 1.24 | 1.23 | 0.88 | 0.87 | 42.97 | 43.31 | 72.79 | 72.89 |
| 5 | -1 | 0 | -1 | 0 | 1.20 | 1.18 | 0.84 | 0.83 | 41.21 | 40.33 | 70.2 | 70.16 |
| 6 | 0 | 0 | 0 | 0 | 1.39 | 1.36 | 1.00 | 1.01 | 50.87 | 49.99 | 76.49 | 76.23 |
| 7 | 1 | 0 | 0 | 1 | 1.10 | 1.13 | 0.78 | 0.81 | 37.02 | 38.12 | 67.32 | 66.87 |
| 8 | 1 | 0 | -1 | 0 | 1.17 | 1.13 | 0.84 | 0.81 | 40.08 | 38.23 | 69.11 | 67.98 |
| 9 | 0 | 1 | 1 | 0 | 1.34 | 1.32 | 1.00 | 1.01 | 50.8 | 49.55 | 73.49 | 72.12 |
| 10 | 0 | -1 | -1 | 0 | 1.33 | 1.37 | 0.96 | 0.97 | 50.06 | 52.75 | 74.42 | 75.44 |
| 11 | 0 | -1 | 0 | -1 | 1.38 | 1.36 | 0.99 | 0.96 | 51.57 | 50.88 | 76.37 | 74.18 |
| 12 | 1 | 0 | 0 | -1 | 1.25 | 1.25 | 0.90 | 0.90 | 43.97 | 45.01 | 71.69 | 73.23 |
| 13 | 0 | 1 | -1 | 0 | 1.13 | 1.14 | 0.80 | 0.81 | 38.65 | 39.44 | 68.62 | 68.55 |
| 14 | 0 | 0 | 0 | 0 | 1.39 | 1.40 | 1.00 | 1.02 | 50.87 | 50.92 | 76.49 | 76.12 |
| 15 | -1 | 0 | 0 | 1 | 1.40 | 1.42 | 0.99 | 1.01 | 51.2 | 51.6 | 77.42 | 75.53 |
| 16 | 0 | -1 | 0 | 1 | 1.31 | 1.26 | 0.93 | 0.91 | 48.3 | 46.11 | 75.20 | 74.46 |
| 17 | -1 | 0 | 1 | 0 | 1.41 | 1.41 | 1.02 | 1.02 | 52.28 | 52.11 | 77.01 | 77.56 |
| 18 | 0 | 0 | -1 | -1 | 1.3 | 1.31 | 0.91 | 0.94 | 44.3 | 43.21 | 72.94 | 72.78 |
| 19 | 0 | -1 | 1 | 0 | 1.33 | 1.34 | 0.94 | 0.95 | 49.13 | 49.78 | 74.15 | 73.87 |
| 20 | -1 | -1 | 0 | 0 | 1.29 | 1.29 | 0.91 | 0.92 | 47.68 | 47.31 | 73.14 | 74.41 |
| 21 | 0 | 0 | 1 | 1 | 1.41 | 1.42 | 1.00 | 0.98 | 50.56 | 52.23 | 75.36 | 76.45 |
| 22 | 1 | 1 | 0 | 0 | 1.06 | 1.08 | 0.77 | 0.78 | 36.23 | 37.18 | 64.3 | 63.96 |
| 23 | 0 | 0 | 0 | 0 | 1.39 | 1.41 | 1.00 | 0.98 | 50.87 | 51.70 | 76.49 | 77.12 |
| 24 | 0 | 1 | 0 | 1 | 1.29 | 1.26 | 0.94 | 0.93 | 47.34 | 46.01 | 73.27 | 74.88 |
| 25 | 0 | 0 | 1 | -1 | 1.38 | 1.39 | 1.01 | 1.03 | 51.86 | 52.10 | 76.34 | 76.90 |
| 26 | -1 | 1 | 0 | 0 | 1.23 | 1.24 | 0.89 | 0.89 | 44.00 | 44.67 | 73.01 | 73.02 |
| 27 | 1 | 0 | 1 | 0 | 1.16 | 1.14 | 0.83 | 0.81 | 40.24 | 39.10 | 66.90 | 66.36 |

### 3.1 Impact of experimental conditions on total phenolic content and total flavonoid content

Table 3 reveals the results of ANOVA for the models which were estimated to investigate the TPC and TFC responses. The model F-values of TPC and TFC were found to be 19.07 and 18.48 respectively; and the *p*-values were obtained lower than 0.05, confirming the significance of the model terms. The main factors like A, B, C and quadratic coefficients including A^2^, B^2^ and the interaction coefficients like AC, AD, BC, BD were found to be significant in the models designed for TPC and TFC both with *p* values < 0.0001.

**Table 3:** Analysis of variance for response variables Y1 (TPC) and Y2 (TFC) recorded during the experimental runs. Significant was assumed at *p<0.05*.

|  | TPC (mg GAE/g extract) |  |  |  |  | TFC (mg QE/g extract) |  |  |  |  |
| --- | --- | --- | --- | --- | --- | --- | --- | --- | --- | --- |
| Source | Sum of squares | Degree of freedom | Mean square | F-value | p-value | Sum of squares | Degree of freedom | Mean square | F-value | p-value |
| Model | 0.278 | 12 | 0.023 | 19.07 | <.0001*** | 0.156 | 12 | 0.013 | 18.4800 | <.0001*** |
| A-Temp. | 0.059 | 1 | 0.059 | 59.0049 | <.0001*** | 0.027 | 1 | 0.027 | 46.9170 | <.0001*** |
| B-Time | 0.027 | 1 | 0.027 | 27.1693 | 0.0002*** | 0.007 | 1 | 0.007 | 12.1444 | 0.0045** |
| C- Extr. Vol. | 0.030 | 1 | 0.030 | 30.1045 | 0.0001*** | 0.023 | 1 | 0.023 | 39.0469 | <.0001*** |
| D-MeOH | 0.000 | 1 | 0.000 | 0.0753 | 0.7884 | 0.000 | 1 | 0.000 | 0.0000 | 1 |
| A<sup>2</sup> | 0.064 | 1 | 0.064 | 63.8134 | <.0001*** | 0.039 | 1 | 0.039 | 67.0193 | <.0001*** |
| B<sup>2</sup> | 0.026 | 1 | 0.026 | 26.5289 | <.0001*** | 0.011 | 1 | 0.011 | 19.6409 | .0008** |
| C<sup>2</sup> | 0.003 | 1 | 0.003 | 3.4761 | 0.0868 | 0.002 | 1 | 0.002 | 3.2860 | 0.0949 |
| D<sup>2</sup> | 0.002 | 1 | 0.002 | 1.9661 | 0.1861 | 0.002 | 1 | 0.002 | 3.3951 | 0.0902 |
| AB | 0.001 | 1 | 0.001 | 1.2293 | 0.2892 | 0.001 | 1 | 0.001 | 1.0830 | 0.3185 |
| AC | 0.012 | 1 | 0.012 | 12.1422 | 0.0045** | 0.009 | 1 | 0.009 | 15.6390 | 0.0019** |
| AD | 0.024 | 1 | 0.024 | 24.1087 | <.0001*** | 0.013 | 1 | 0.013 | 22.9170 | 0.0004*** |
| BC | 0.011 | 1 | 0.011 | 11.0634 | <.0001*** | 0.012 | 1 | 0.012 | 20.9675 | 0.0006*** |
| BD | 0.006 | 1 | 0.006 | 5.6446 | 0.0350* | 0.004 | 1 | 0.004 | 6.2383 | 0.0280* |
| CD | 0.001 | 1 | 0.001 | 0.6272 | 0.4437 | 0.000 | 1 | 0.000 | 0.3899 | 0.5440 |
| Residual | 0.012 | 12 | 0.001 |  |  | 0.007 | 12 | 0.001 |  |  |
| Lack of fit | 0.011 | 10 | 0.001 | 1.5083 | 0.4634 | 0.006 | 10 | 0.001 | 1.3981 | 0.4875 |
| IPure error | 0.001 | 2 | 0.001 |  |  | 0.001 | 2 | 0.000 |  |  |
| R<sup>2</sup> | 0.957 |  |  |  |  | 0.9557 |  |  |  |  |
| Adjusted R<sup>2</sup> | 0.9068 |  |  |  |  | 0.9039 |  |  |  |  |
| C.V.% | 7.90 |  |  |  |  | 8.25 |  |  |  |  |

**Table 4.** Analysis of variance for the response variables Y3 (%DPPH) and Y4 (%ABTS) recorded during the experimental runs. Significant was assumed at *p<0.05*.

|  | %DPPH |  |  |  |  | %ABTS |  |  |  |  |
| --- | --- | --- | --- | --- | --- | --- | --- | --- | --- | --- |
| Source | Sum of squares | Degree of freedom | Mean square | F-value | p-value | Sum of squares | Degree of freedom | Mean square | F-value | p-value |
| Model | 681.331 | 12 | 56.778 | 17.32 | <.0001*** | 329.853 | 12 | 27.488 | 11 | <.0001*** |
| A- Temp. | 130.021 | 1 | 130.021 | 48.5532 | <.0001*** | 94.136 | 1 | 94.136 | 47.3875 | <.0001*** |
| B- Time | 71.151 | 1 | 71.151 | 26.5696 | 0.0002*** | 31.299 | 1 | 31.299 | 15.7556 | 0.0018** |
| C- Extr. Vol. | 94.529 | 1 | 94.529 | 35.2995 | <.0001*** | 15.893 | 1 | 15.893 | 8.0004 | 0.0152* |
| D- MeOH | 1.229 | 1 | 1.229 | 0.4589 | 0.511 | 0.046 | 1 | 0.046 | 0.0230 | 0.882049 |
| A<sup>2</sup> | 170.691 | 1 | 170.691 | 63.7404 | <.0001*** | 70.381 | 1 | 70.381 | 35.4291 | <.0001*** |
| B<sup>2</sup> | 18.293 | 1 | 18.293 | 6.8309 | 0.0226* | 19.125 | 1 | 19.125 | 9.6272 | 0.0091** |
| C<sup>2</sup> | 6.584 | 1 | 6.584 | 2.4585 | 0.1428 | 15.293 | 1 | 15.293 | 7.6986 | 0.0168* |
| D<sup>2</sup> | 6.049 | 1 | 6.049 | 2.2589 | 0.1586 | 0.109 | 1 | 0.109 | 0.0548 | 0.8187 |
| AB | 1.404 | 1 | 1.404 | 0.5244 | 0.4828 | 9.641 | 1 | 9.641 | 4.8532 | 0.0478* |
| AC | 29.757 | 1 | 29.757 | 11.1121 | 0.0059** | 20.340 | 1 | 20.34 | 10.2391 | 0.0076** |
| AD | 57.608 | 1 | 57.608 | 21.5124 | 0.0005*** | 20.250 | 1 | 20.25 | 10.1937 | 0.0077** |
| BC | 42.772 | 1 | 42.772 | 15.9721 | 0.0017** | 6.605 | 1 | 6.605 | 3.3249 | 0.0932 |
| BD | 15.327 | 1 | 15.327 | 5.7236 | 0.0339* | 1.677 | 1 | 1.677 | 0.8442 | 0.3762 |
| CD | 3.783 | 1 | 3.783 | 1.4127 | 0.2576 | 1.221 | 1 | 1.221 | 0.6147 | 0.4482 |
| Residual | 32.135 | 12 | 2.678 |  |  | 23.838 | 12 | 1.987 |  |  |
| Lack of fit | 30.669 | 10 | 3.067 | 4.1846 | 0.2081 | 23.237 | 10 | 2.324 | 7.7276 | 0.1199 |
| Pure error | 1.466 | 2 | 0.733 |  |  | 0.601 | 2 | 0.301 |  |  |
| R<sup>2</sup> | 0.9528 |  |  |  |  | 0.9277 |  |  |  |  |
| Adjusted R<sup>2</sup> | 0.8978 |  |  |  |  | 0.8434 |  |  |  |  |
| C.V.% | 10.84 |  |  |  |  | 4.70 |  |  |  |  |

Furthermore, the high values of R^2^ (0.96 for TPC and TFC both) and Adj-R^2^ (0.90 for both TPC and TFC), the coefficient of variation:CV (7.90 and 8.25 for TPC and TFC respectively) and insignificant values of lack of fit (*p* > 0.05, TPC: 0.46; TFC: 0.48) validated the equations of the numerical model and indicated the ability for prediction of the total phenolic content and total flavonoid content in accordance with the different combinational values of the independent variables.

In addition, accuracy of the regression models were examined by assessing the diagnostic plots of predicted versus actual values. Figure 1 demonstrates the comparative response of predicted and actual values and authenticates that the experimental values are in close proximity of the predicted values, suggesting an excellent match with no significant deviation.

**Figure 1:**
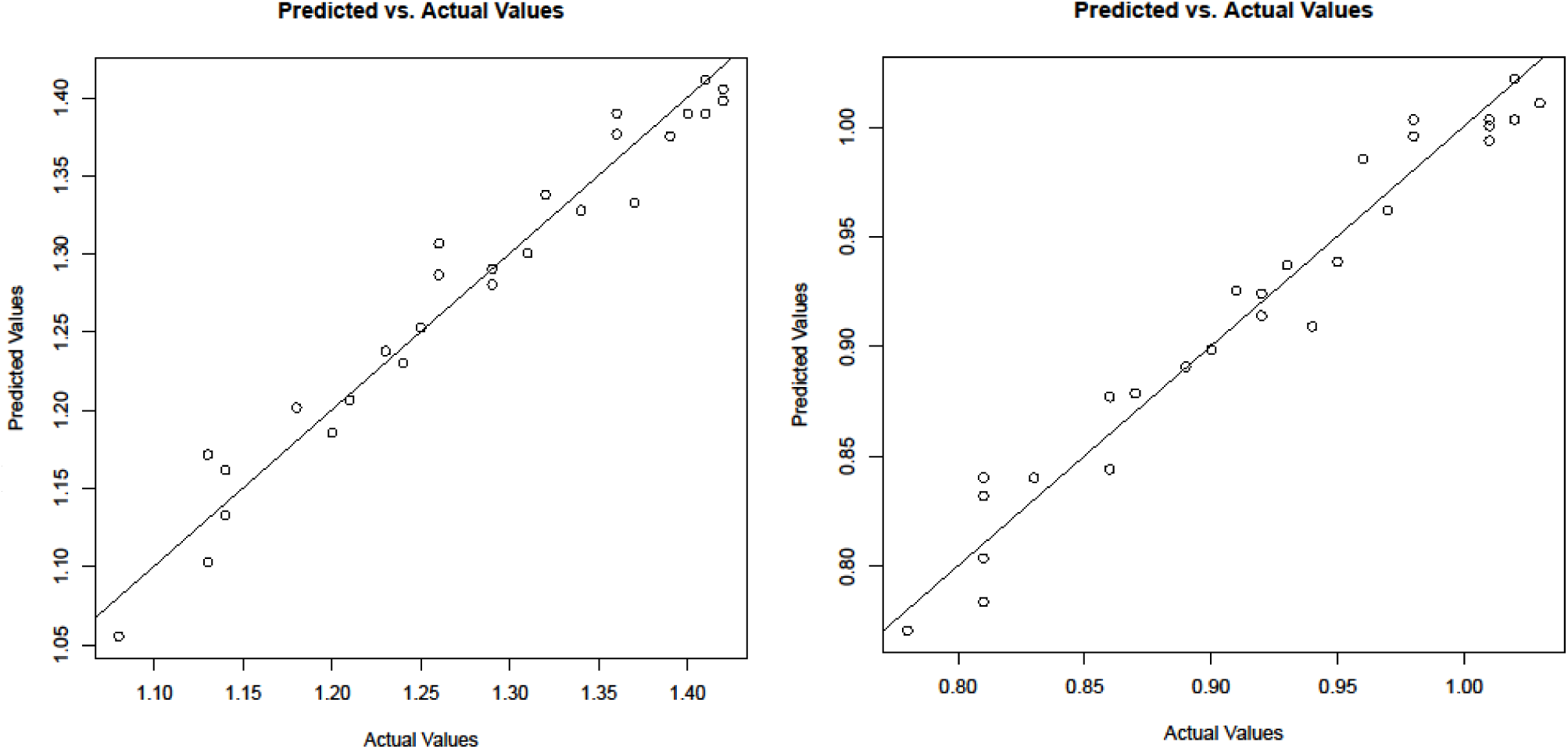
Diagnostic plots corresponding to the estimated models for predicting TPC and TFC.

**Figure 2:**
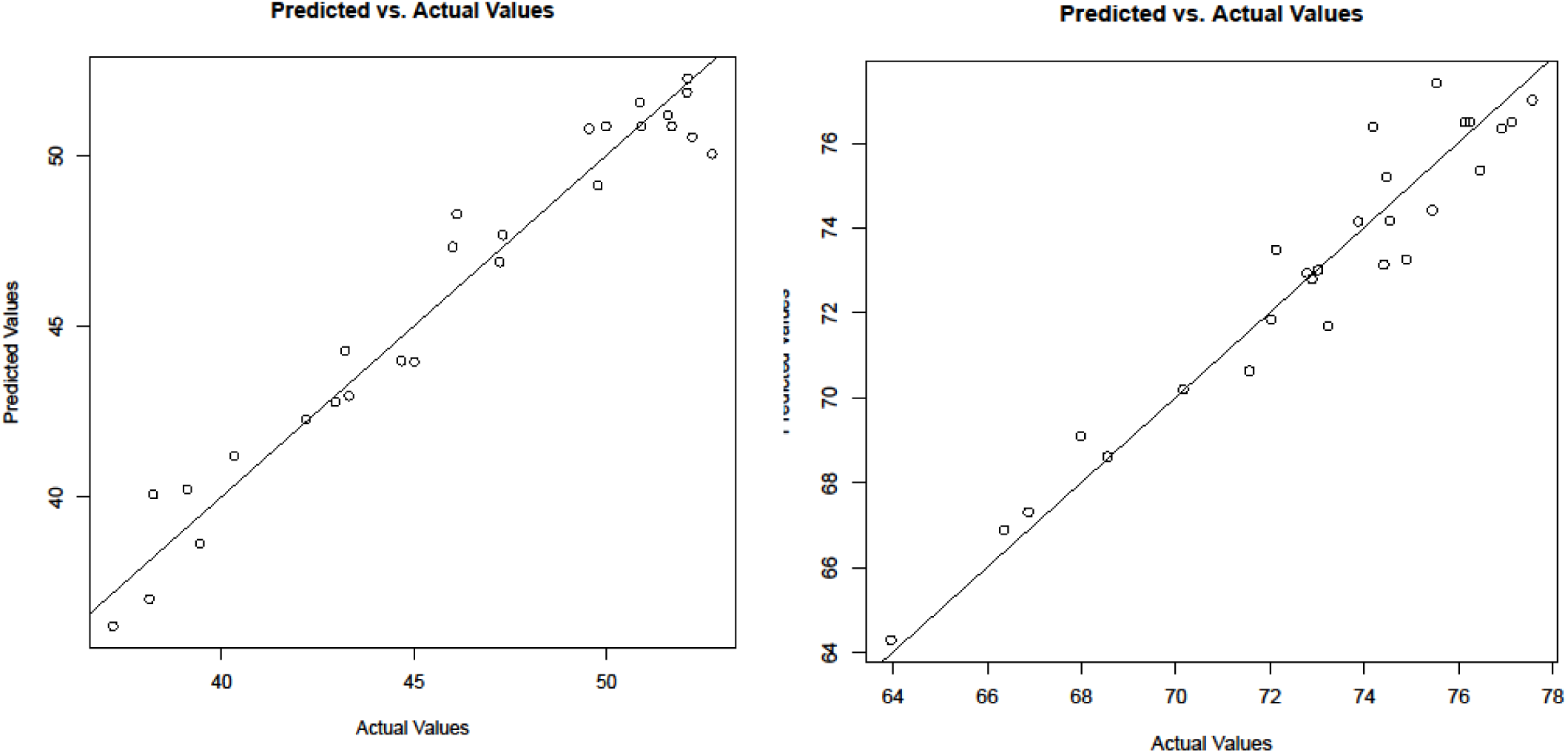
Diagnostic plots corresponding to the estimated models for predicting for DPPH and ABTS.

Figures 3 and 4 present the response surface plots to understand the influence of various independent variables on TPC and TFC respectively, which we discuss next. Time and volume are found to be critical factors to influence both of TPC (Figures 3 (c) and 3 (d) for time, and Figures 3 (a) and 3 (c) for volume) and TFC (Figures 4 (c) and 4 (d) for time, and Figure 4 (a) and 4 (c) for volume). TPC and TFC increases with the increase of time up to a certain point and start decreasing beyond that. TPC and TFC shows increasing trend with volume. Usually, increased time and extraction volume can boost the solubility of bioactive compounds like phenolic and flavonoid contents from the plant matrices and allowing their diffusion into the extraction solvent [35]. Our research findings also support the earlier report where the study claimed that the increased amount of extraction solvent is responsible for high permeability to the plant matrices resulting swelling of the cell wall and cell membrane [27], [36]. This mechanism increases the penetration of the extraction solvent towards the plant matrices and causing the significant interaction with the extraction solvent and the phytoconstituents [37]. It is well-known that the polarity index of the extraction solvent is having a positive influence with the extraction of the polar bioactive compounds like phenolics and flavonoids. So, the higher solvent volume can be responsible for the increased amounts of extractions of these polar compounds [38].

**Figure 3:**
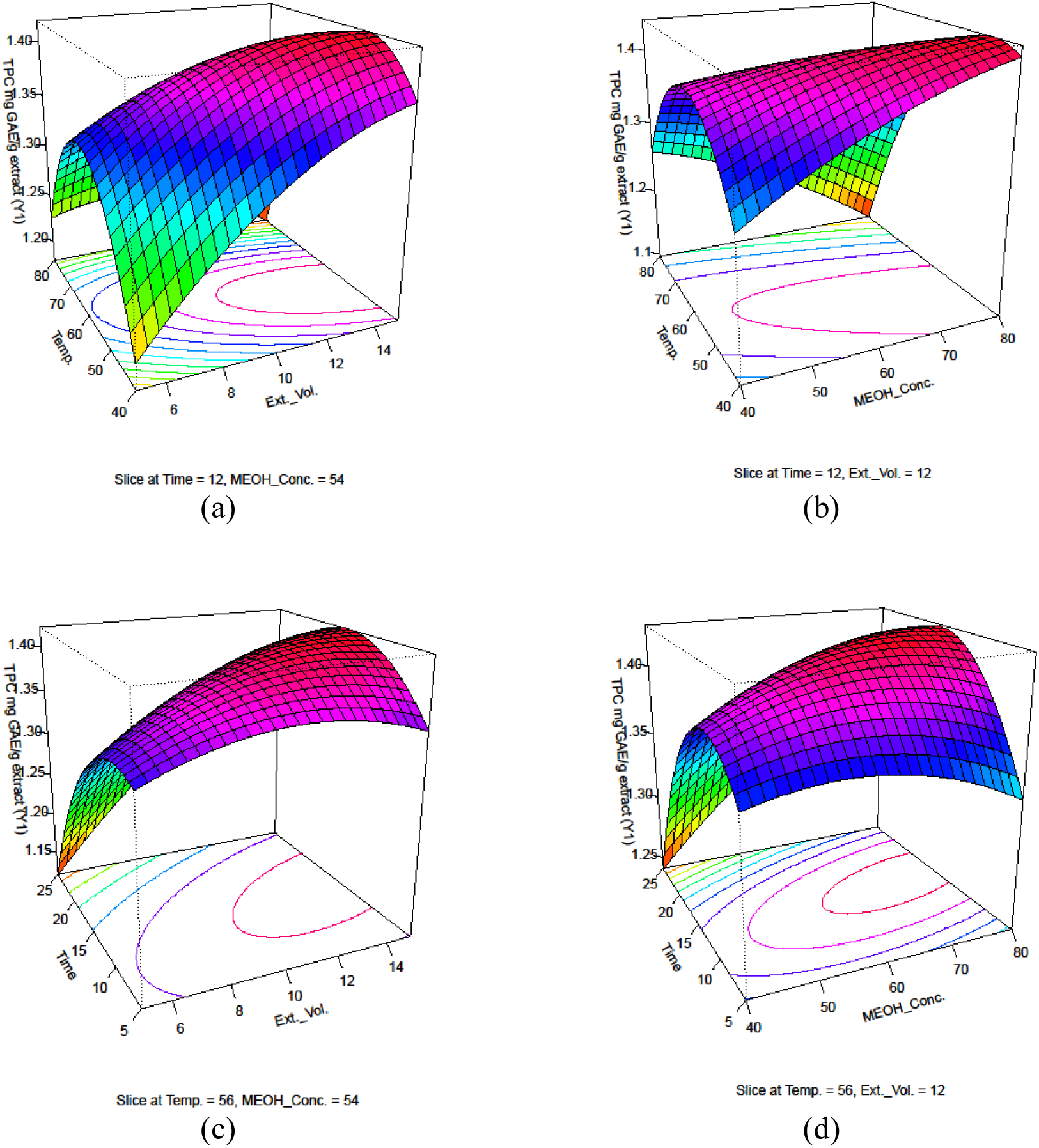
Response surface plots depicting significant interactions between two independent variables on TPC (Y1): (a) temperature vs extraction volume; (b) temperature vs MEOH concentration; (c) time vs extraction volume; (d) time vs MEOH concentration

**Figure 4:**
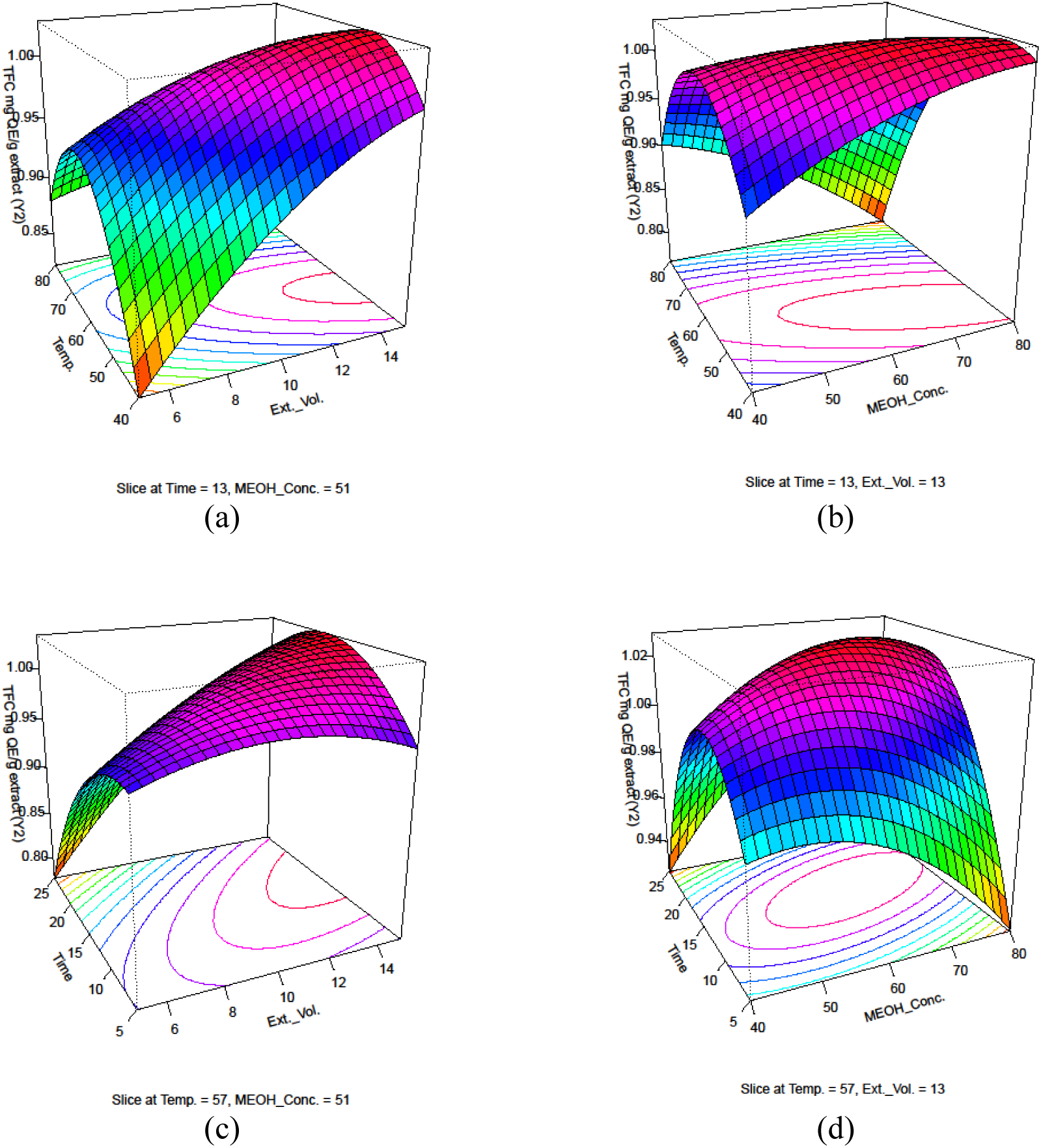
Response surface plots depicting significant interactions between two independent variables on TFC (Y2): (a) temperature vs extraction volume; (b) temperature vs MEOH concentration; (c) time vs extraction volume; (d) time vs MEOH concentration

For MAE, the extraction time has a critical role on extraction of the bioactive constituents from the plant matrix [39]. Longer duration of exposure to microwave radiation causes more breakage of cellular wall and consequently promotes the higher rate of mass transfer from the sample to extraction solvent leading to enhancement of the total phenolic and total flavonoid contents [40]. But the plant materials can be overheated when the exposure of the microwave radiation is extended over the period. So, to find the ideal time of microwave irradiation, process optimization is required [40], [41], [42].

It is also evident that the extraction of TFC is increasing with the time and the methanol concentration but after an ideal time and concentration, the extraction of the TFC is getting negatively affected (Figure 4 (d)).

Figures 3 (a) and 3 (b) for TPC, and Figures 4 (a) and 4 (b) for TFC show that beyond a certain temperature the response values are getting decreased. This is because of denaturation of the heat sensitive bioactive components at the higher temperature. Kinetic energy can be increased at the elevated temperature might cause the degradation or structural changes of the important specific components, lowering the efficiency of the extraction [43], [44].

Phenolics and flavonoids are the two vital indicators predominantly employed to represent the plant-based bioactivities in leaf extract. The experiments carried out so far have validated that the leaf extracts of *Vitex* possessed ample amounts of phenolics and flavonoid contents. The previous study revealed that the leaf extracts of *Vitex* a wide array of polyphenols (aspartic acid, gallic acid, dihydroxy benzoic acid, catechin, chlorogenic acid, vanillic acid, rutin, trans-cinnamic acid, ferulic acid, quercetin, apigenin, kaempferol, and tannic acid) was detected by HPLC separation with the distinct elution times [16]. For this study, we used BBD to optimize the conditions for the microwave-assisted extraction. Because that research work demonstrated these highly sensitive extraction conditions as certain and robust for bringing out the highest amounts of phenolic compounds from the plant cells without damaging their bioactivities and using a minimum laboratory set up cost as well [30]. Furthermore, Elfalleh et al., (2019) demonstrated the use of different polar solvents as solvents for successful extractions of phenolics and flavonoid compounds [45]. Phenolics and flavonoids both are the most important phyto-constituents which are accounted for by their role in scavenging potential or chelating ability. These phytochemicals are widely known as promising source of natural antioxidants and also have been reported for exhibiting the anticancer, antibacterial, cardioprotective, anti-inflammatory, and immune system promoting activities and which leads to project those compounds as an excellent drug candidate for pharmaceutical field [46].

### 3.2 Impact of experimental conditions on antioxidant activities of DPPH and ABTS

Table 4 displays the results of ANOVA for the RSM models, which were employed to investigate the DPPH and ABTS activities. The model F-values of DPPH and ABTS were found to be 17.32 and 11 respectively; and the *p*-values obtained were lower than the 0.05 confirming the significance of the model terms. The main factors like A (*p* < 0.0001), B (*p* = 0.0002), C (*p* < 0.0001), and quadratic coefficients including A^2^ (*p* < 0.0001), B^2^ (*p* = 0.0226), and the interaction coefficients like AC (*p* = 0.0059), AD (*p* = 0.0005), BC (*p =* 0.0017), BD (*p* = 0.0339) were found to be significant in the model designed for DPPH. For the model of ABTS, the main factors including A (*p* < 0.0001), B (*p* = 0.0018), C (*p* = 0.0152), and quadratic factors like A^2^ (*p* < 0.0001), B^2^ (*p* = 0.0091), C^2^ (*p* = 0.0168), and the interaction coefficients such as AB (*p* = 0.0478), AC (*p* = 0.0076), AD (*p* = 0.0077) were found to be significant.

**Table 4:** Predicted and experimental values of dependent variables at optimum experimental conditions. *Mean values ± standard deviation (*n* = 3). Significant was assumed at *p < 0.05*.

|  | Independent variable |  |  |  |
| --- | --- | --- | --- | --- |
|  | Temperature (°C) | Time (min) | Extr. Volume (mL) | MeOH (%v/v) |
| Optimal conditions | 48.141 | 14.098 | 14.762 | 65.429 |
| Responses | TPC (mg GAE/g extract) | TFC (mg GAE/g extract) | %DPPH | %ABTS |
| Predicted values | 1.452 | 1.046 | 53.856 | 77.752 |

Additionally, the high values of R^2^ (0.95 and 0.93 for DPPH and ABTS respectively) and Adj-R^2^ (0.89 and 0.84 for DPPH and ABTS respectively), the coefficient of variation: CV (10.84 and 4.70 for DPPH and ABTS respectively) and insignificant values of lack of fit (*p* > 0.05, DPPH: 0.21; ABTS: 0.12) validated the equations of the numerical models and indicated the ability for prediction of the antioxidant activities of the DPPH and ABTS in accordance with the different combinational values of the independent variables.

Furthermore, accuracy of the regression models was examined by assessing the Diagnostic plot of predicted versus actual values. Figure 2 demonstrates the comparative response of predicted and actual values and authenticates that the experimental values are in close proximity of the predicted values, suggesting an excellent match with no significant deviation.

Figures 5 and 6 present the response surface plots to understand the influence of various independent variables on DPPH and ABTS potentials respectively, which we discuss next. Time and volume are found to be critical factors to influence both of DPPH (Figures 5 (c) and 5 (d) for time, and Figures 5 (a) and 5 (c) for volume), and ABTS (Figures 6 (c) for time, and Figure 6 (a) for volume). DPPH and ABTS increases with the increase of time up to a certain point and start decreasing beyond that. DPPH and ABTS shows increasing trend with volume in general.

**Figure 5:**
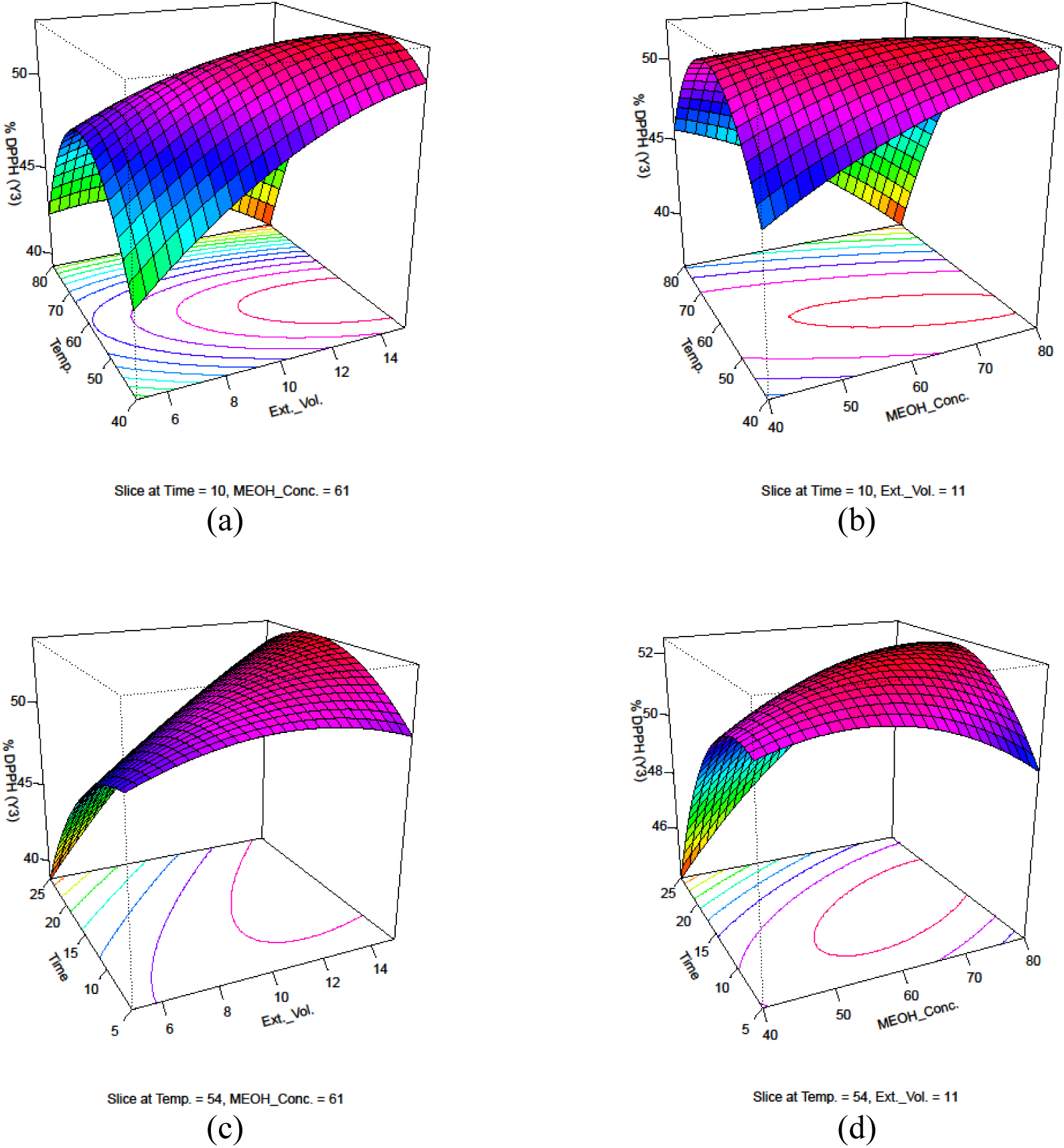
Response surface plots depicting significant interactions between two independent variables on % DPPH (Y3): (a) temperature vs extraction volume; (b) temperature vs MEOH concentration; (c) time vs extraction volume; (d) time vs MEOH concentration

**Figure 6:**
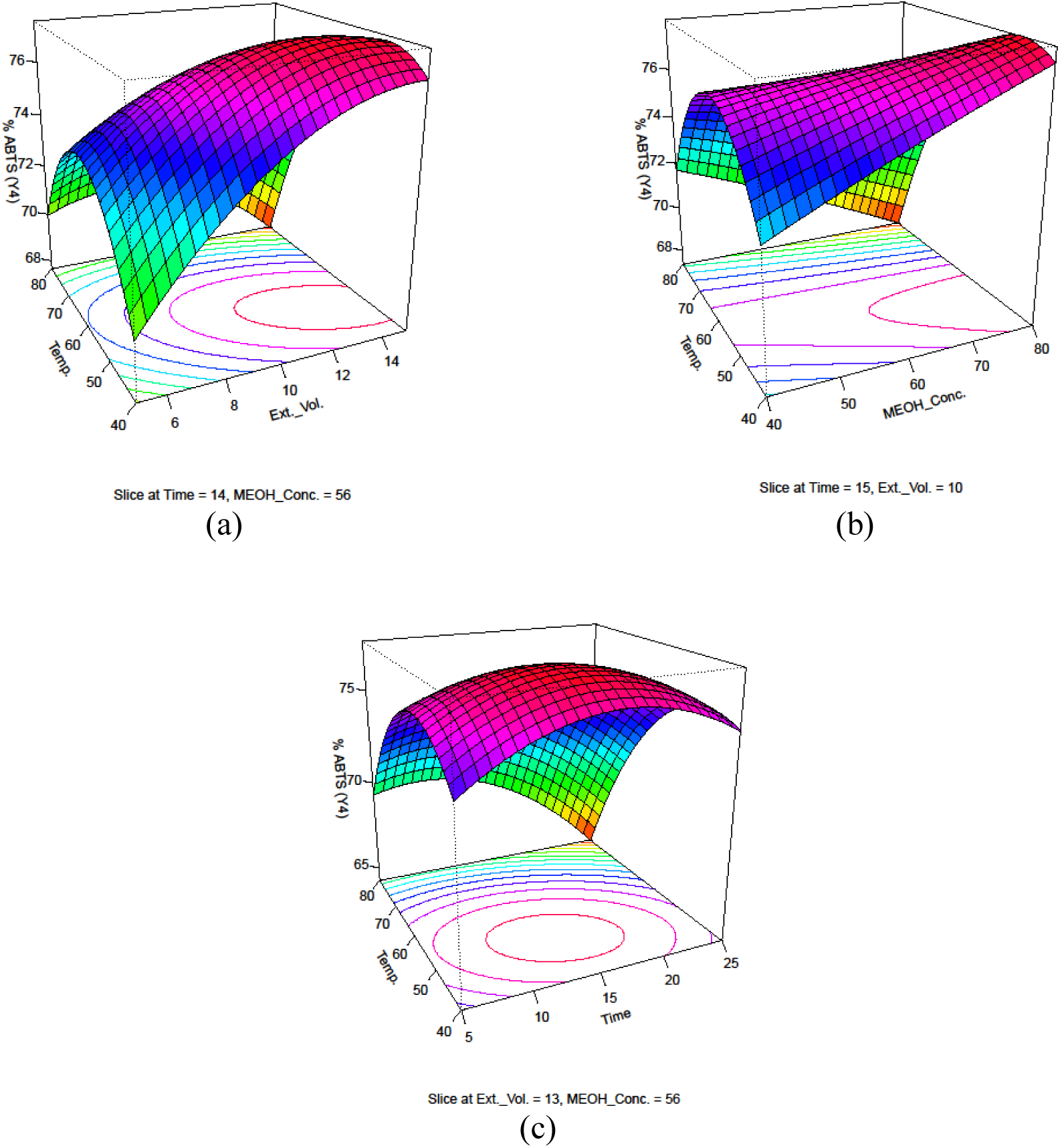
Response surface plots depicting significant interactions between two independent variables on % ABTS (Y4): (a) temperature vs time; (b) temperature vs extraction volume; (c) temperature vs MEOH concentration

Figures 5 (a) and 5 (b) for DPPH, and Figures 6 (a) and 6 (b) for ABTS show that the after increment of a certain temperature, the response values decrease. Similar trend is observed for the influence of methanol concentration on the antioxidant activities of DPPH (Figures 5 (b) and 5 (d)), but for ABTS, the effect of methanol concentration in inconclusive (Figures 6 (b)). It is already discussed that higher temperature is associate with the deterioration of some bioactive compounds and less efficient extraction which leads to decrease the antioxidant potentials of DPPH and ABTS [47]. Extraction efficiency also depends on the polarity of the solvent and increased solvent volume is also responsible for the more efficient extraction of the polar compounds. But it was reported that the elevated concentration of ethanol could destroy the protein structure and the hydration level of plant cell, eventually hampers the extraction efficiency and lowers the values of yields [48].

*In-vitro* antioxidant assays were predominantly executed to build up a direct relationship between the degree of scavenging activity and therapeutic applications[49]. To our knowledge, the rate of electron transfer of ABTS is much quicker than DPPH; and this kinetic reaction makes a subtle difference in mechanism of actions of DPPH and ABTS at antioxidant activity level. DPPH radical reaction is worked on the use of very crowded radical whereas the ABTS is applicable for low crowded radical. According to Mareček et al. 2017 the DPPH radical gets chemically involved with polyphenols (catechins, proanthocyanidins), and the highly reactive ABTS gets chemically involved with a broader range of antioxidants [50]. Here, the leaf extract demonstrated a remarkable antioxidant potential against the highly reactive species like hydroxyl radical, singlet oxygen, non-radical reactive oxygen species. Consequently, the leaf extract may play an important role in mitigating cellular stress which in turn plays a profound role in arresting a wide range of onset diseases such as diabetes, cancer, cardiovascular diseases, eye disorders, arthritis, obesity, autoimmune diseases, inflammatory bowel disease [51].

### 3.3 Simultaneous multi-response optimization

Optimum experimental conditions for MAE were obtained by simultaneously optimizing the four response variables using desirability function approach [52]. The mathematical optimization model was used to investigate the optimum levels for temperature, extraction time, extraction volume and methanol concentration, with a target of highest possible values of the dependent variables. As per the RSM, the optimal conditions for MAE were identified as a temperature of 48°C for 14 min, utilizing 15 mL of 65% (v/v) methanol. The quadratic model has a *p*-value of < 0.0001, *p*-value of the lack of fit of 0.4634, R^2^: 0.957 and adjusted R^2^: 0.9068. To authenticate the values of predicted dependent variables, the extraction was carried out three times at the obtained optimal conditions. The predicted and experimental values of the four response variables are shown in Table 4. The experimental values found for TPC, TFC and antioxidant assays (DPPH free radical scavenging assay, ABTS radical scavenging assay) were in close proximity with the results predicted by the polynomial quadratic models, showing credibility and effectiveness of the optimized MAE-BBD-RSM model. This technique reduces the number of runs of the experiment necessary for the polyphenol extraction with their antioxidant potentials with maintaining the credibility and effectiveness of the results.

### 3.4 GC-MS analysis

The qualitative analysis of leaf extract was investigated by GC-MS. Total 15 peaks were identified by comparing the percentage area of peak, retention time and the pattern of mass spectral fragmentation of the widely known compounds those are existed in the database of NIST. Among 15 peaks, 12 identified bioactive phytochemical constitutes of methanol extract is summarized in Figure 7 and Table 5 respectively, along with their names, molecular formulas, molecular weight and class.

**Figure 7:**
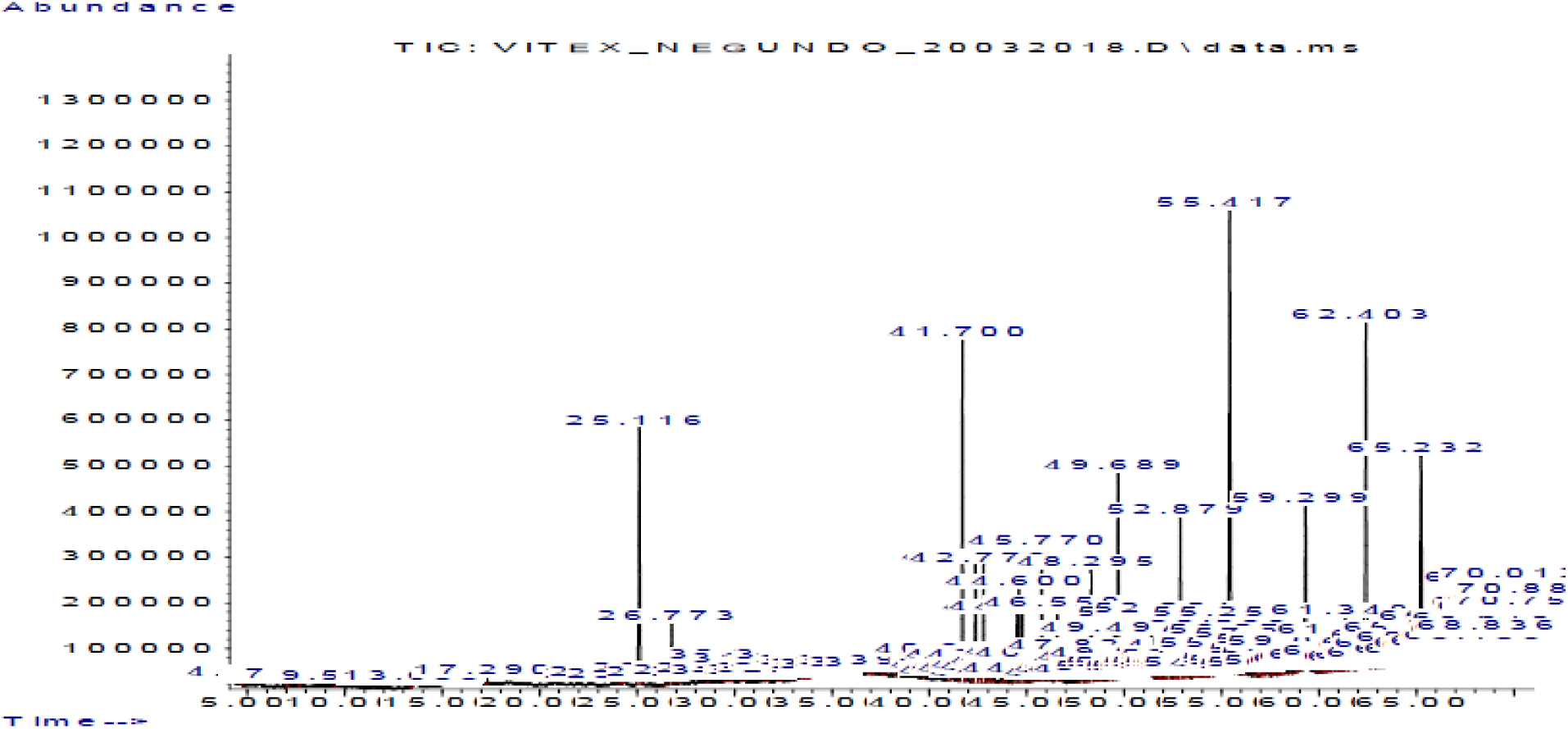
Chromatograms of volatile organic compounds were obtained from the methanol extract of *Vitex negundo* leaf.

**Table 5:** The identified volatile components of methanol extract of *Vitex negundo* leaf are enlisted.

| Retention time (min) | % Area of peak | Compound identified | Molecular formula | Molecular weight | Class of compound |
| --- | --- | --- | --- | --- | --- |
| 24.048 | 0.05 | Lactose | C<sub>12</sub>H<sub>22</sub>O<sub>11</sub> | 342.3 | Disaccharide |
| 25.116 | 4.72 | α-caryophyllene | C<sub>15</sub>H<sub>24</sub> | 204.35 | Sesquiterpenoids |
| 26.772 | 1.05 | β-myrcene | C<sub>10</sub>H<sub>16</sub> | 136.23 | Monoterpene |
| 38.358 | 0.04 | 1-methylethyl ester hexanoic acid | C<sub>9</sub>H<sub>18</sub>O<sub>2</sub> | 158.24 | Fatty acid ester |
| 43.116 | 0.26 | α-farnesene | C<sub>15</sub>H<sub>24</sub> | 204.35 | Sesquiterpene |
| 43.904 | 0.10 | 3-methyl-decanoic acid | C<sub>11</sub>H<sub>22</sub>O<sub>2</sub> | 186.29 | Fatty acid |
| 45.570 | 0.38 | Thunbergol | C<sub>20</sub>H<sub>34</sub>O | 290.5 | Monocyclic diterpene alcohol |
| 48.709 | 0.20 | (Isomer 1) cis-trans citral | C<sub>10</sub>H<sub>16</sub>O | 152.23 | Monoterpene aldehyde |
| 49.690 | 4.31 | Phytol | C<sub>20</sub>H<sub>40</sub>O | 296.539 | Diterpene alcohol |
| 51.802 | 0.06 | Oleic acid | C<sub>18</sub>H<sub>34</sub>O<sub>2</sub> | 282.5 | Fatty acid |
| 58.049 | 0.03 | 2-myristynoyl-glycinamide | C<sub>16</sub>H<sub>28</sub>N<sub>2</sub>O<sub>2</sub> | 280.41 | Amino compound |
| 61.349 | 1.59 | Squalene | C<sub>30</sub>H<sub>50</sub> | 410.730 | Triterpene |

In recent research trends, the plant extracts have received widespread recognition as the antibacterial, antifungal, antioxidant, insecticidal and flavoring agents. Several research works have approved the critical role of bioactive phytoconstituents in prevention of cell proliferation and inflammatory diseases [15], [53], [54], [55], [56]. Some of the important components of volatile oil such as oleic acid, isocaryophyllene, caryophyllene, phytol, citral have been cited for their promising anti-inflammatory and chemo-preventive property. Presence of monounsaturated fatty acid like oleic acid facilitates to improve the body immune system by successfully eliminating some of pathogens, potentiating the role of macrophages, lymphocytes and neutrophils. Furthermore, β-myrcene, 1-methylethyl ester hexanoic acid, thunbergol, 2-myristynoyl-glycinamide, squalene, γ-sitosterol and asarone have been reported for their anti-inflammatory activity, anticancer activity; antibacterial activity, antifungal activity; antibacterial activity, antioxidant activity; antimicrobial activity; antibacterial activity, antioxidant activity, antitumor activity, cancer preventive activity; antibacterial activity; anticancer activity, antioxidant activity, Anti-alzheimer’s activity [14], [46], [57], [58], [59].

### 3.5 Analysis of tocopherol

Result revealed the presence of tocopherol in methanol extract of *Vitex negundo* leaves (Figure 8). The content of tocopherol in methanol extract was obtained to be 414.87 µg/g. Such results indicated that the leaf of *Vitex negundo* is a very good natural source of tocopherol.

**Figure 8:**
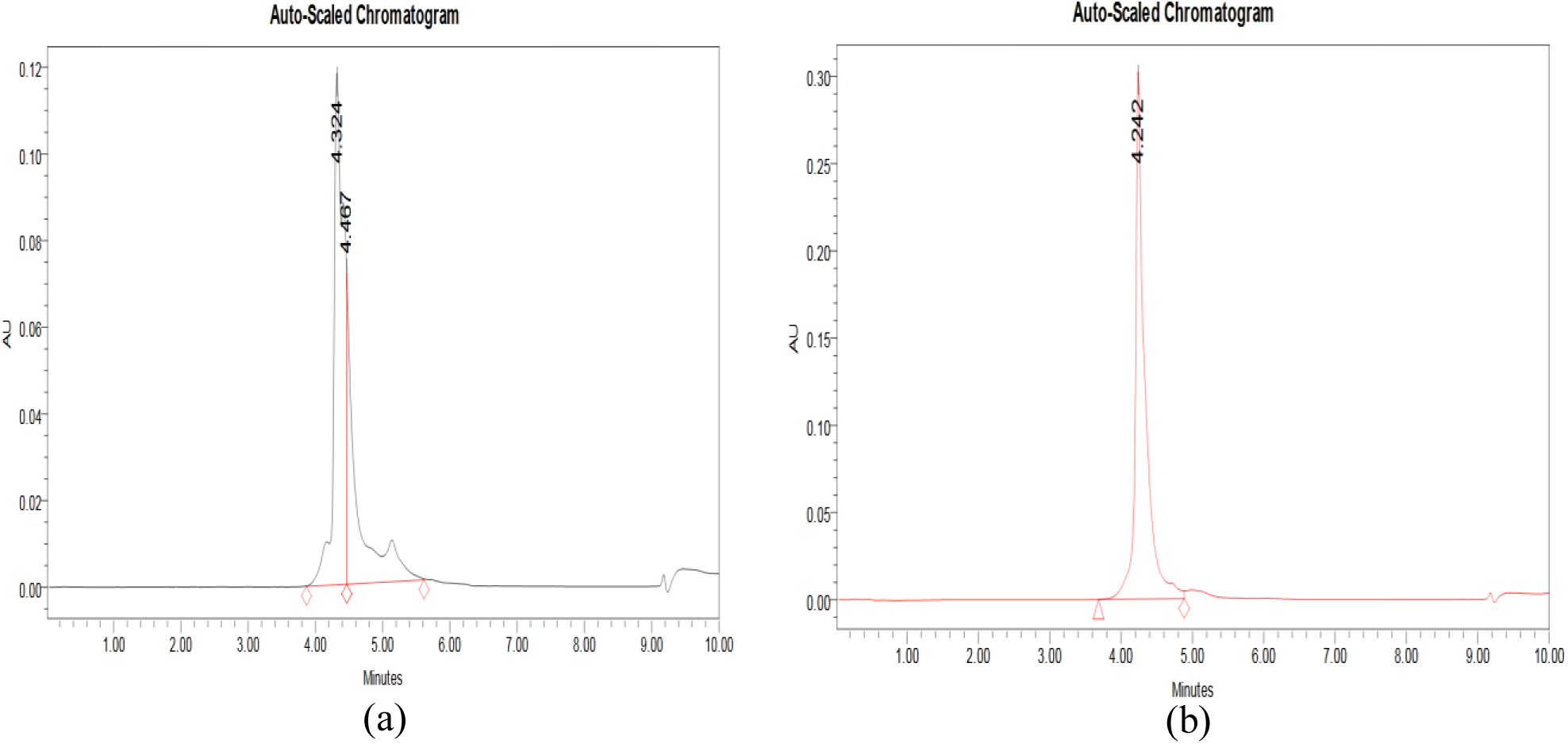
Reversed phase-high-performance liquid chromatography of tocopherol was detected in (a) methanol extract of *Vitex negundo* leaf and (b) standard compound

Tocopherol and its isomers are reported for their strong antioxidant, anti-inflammatory, anti-atherogenic and anticancer activities [60]. Publication records [61], [62], [63] revealed that the polyphenols have been extensively investigated for their several potential clinical significances including antioxidant, antimicrobial, anti-inflammatory, anticancer properties. Furthermore, polyphenols could reduce the detrimental impact on cellular components by promoting the effect of other antibiotics which can act as reactive species scavengers or lipoxygenase inhibitor [64].

### 3.6 Cytotoxic effects of leaf extracts of *Vitex negundo*

To investigate the inhibitory effects on cell viability of human prostate cancer cell lines, MTT assay was carried out by exposing cultured PC_3_ cells to different doses (12.5 μg/mL, 25 μg/mL, 50 μg/mL, 75 µg/mL and 100 μg/mL) of methanol extract for 24 h and 48 h respectively. Figure 11 demonstrates the significant preventive effects of methanol extract on viability of PC_3_ cells in a concentration dependent manner at 24 h. Significant cytotoxic effect was found from the range of doses of 25 μg/mL to 100 μg/mL for 24 h and 48 h both. The 100 μg/mL dose was determined as the strongest significant dose with a value of *p* < 0.0001. Figure 13 clearly indicates the dose dependent significant cytotoxic effects of methanol extract against PC_3_ cell lines at 48 h. Here, the 100 μg/mL concentration was again considered as the strongest effective dose with a value of *p* < 0.0001. The IC_50_ value of methanol extract was found to be 90 μg/mL and 70 μg/mL at 24 h and 48 h respectively.

**Figure 9:**
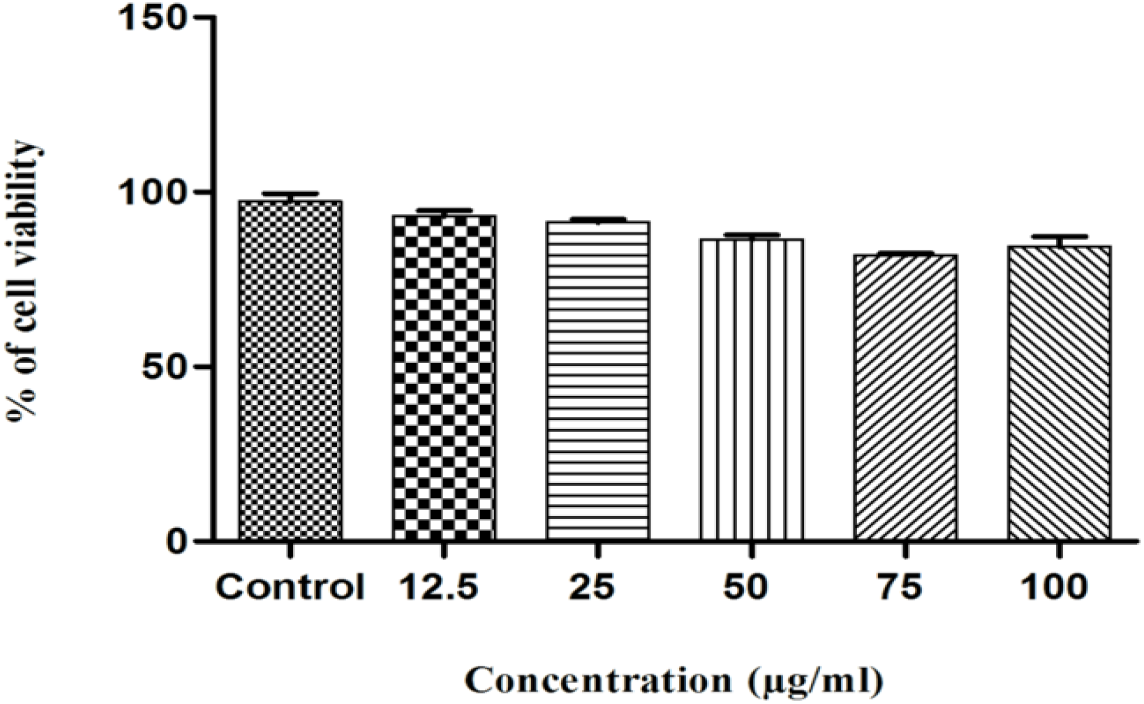
Concentration-dependent cell viability of methanol extract against WI38 cell line at 24 h. The values are expressed as the percentage in comparison with the untreated (control) cell group and each bar in the graph is represented as mean ± SD (n = 3)

**Figure 10:**
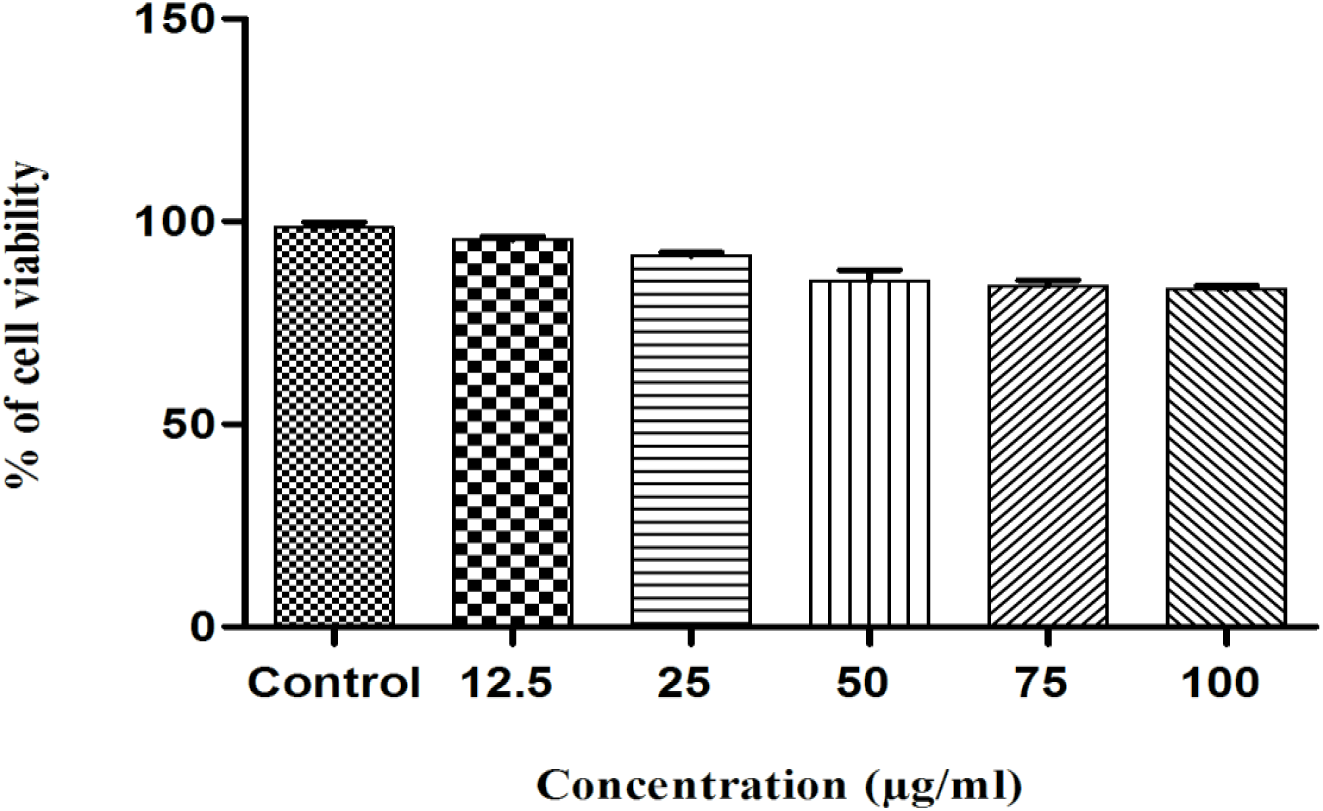
Concentration-dependent cell viability of methanol extract against WI38 cell line at 48 h. The values are expressed as the percentage in comparison with the untreated (control) cell group and each bar in the graph is represented as mean ± SD (n = 3)

**Figure 11:**
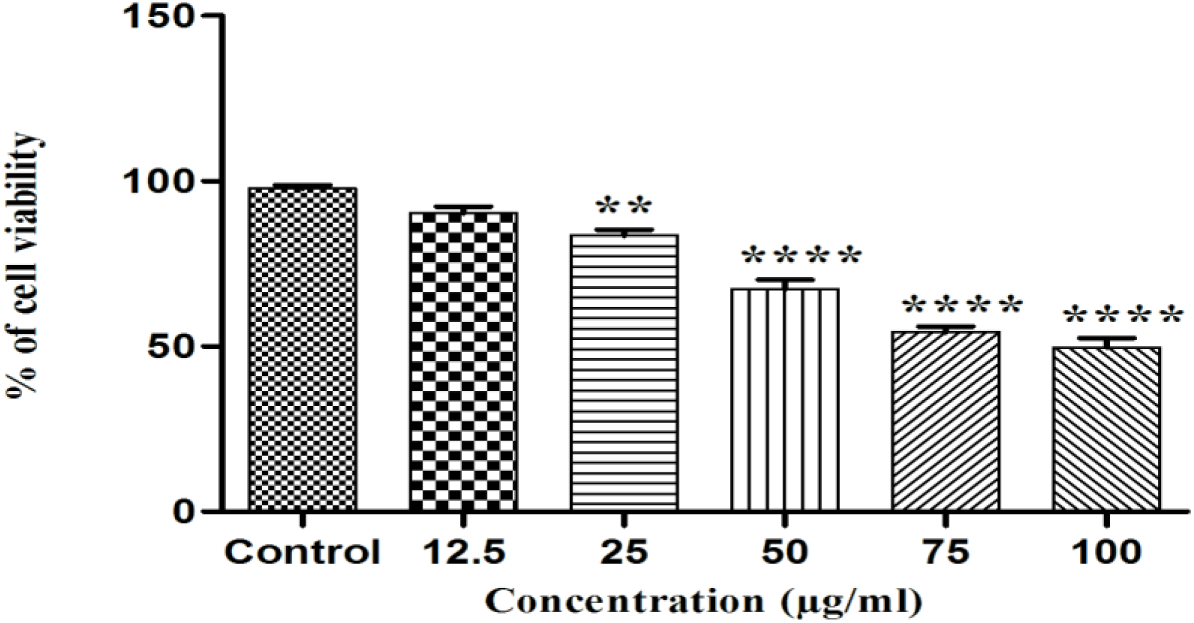
Concentration-dependent cell viability of methanol extract against PC_3_ cell line at 24 h. The PC_3_ cells were treated with leaf extracts (12.5, 25, 50, 75, 100 μg/mL) for 24 h. Results are presented as the mean ± SD of at least triplicates of each experiment. \**p* < 0.05, \*\**p* < 0.01, \*\*\**p* < 0.001 and \*\*\*\**p* < 0.0001, statistically significant in comparison with control groups by one-way ANOVA and followed by Bonferroni’s multiple comparison test

**Figure 12:**
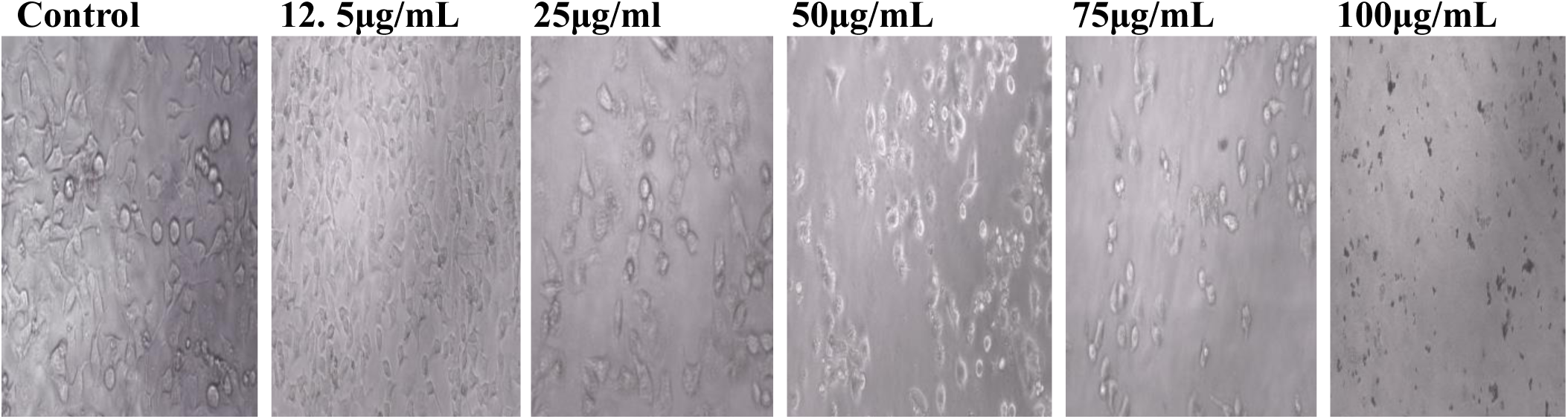
Concentration-dependent finite deformation effects in cell morphology of PC_3_. After treating with the 12.5 μg/mL, 25 μg/mL, 50 μg/mL, 75 μg/mL, 100μg/ml of methanol extract for 24 h

**Figure 13:**
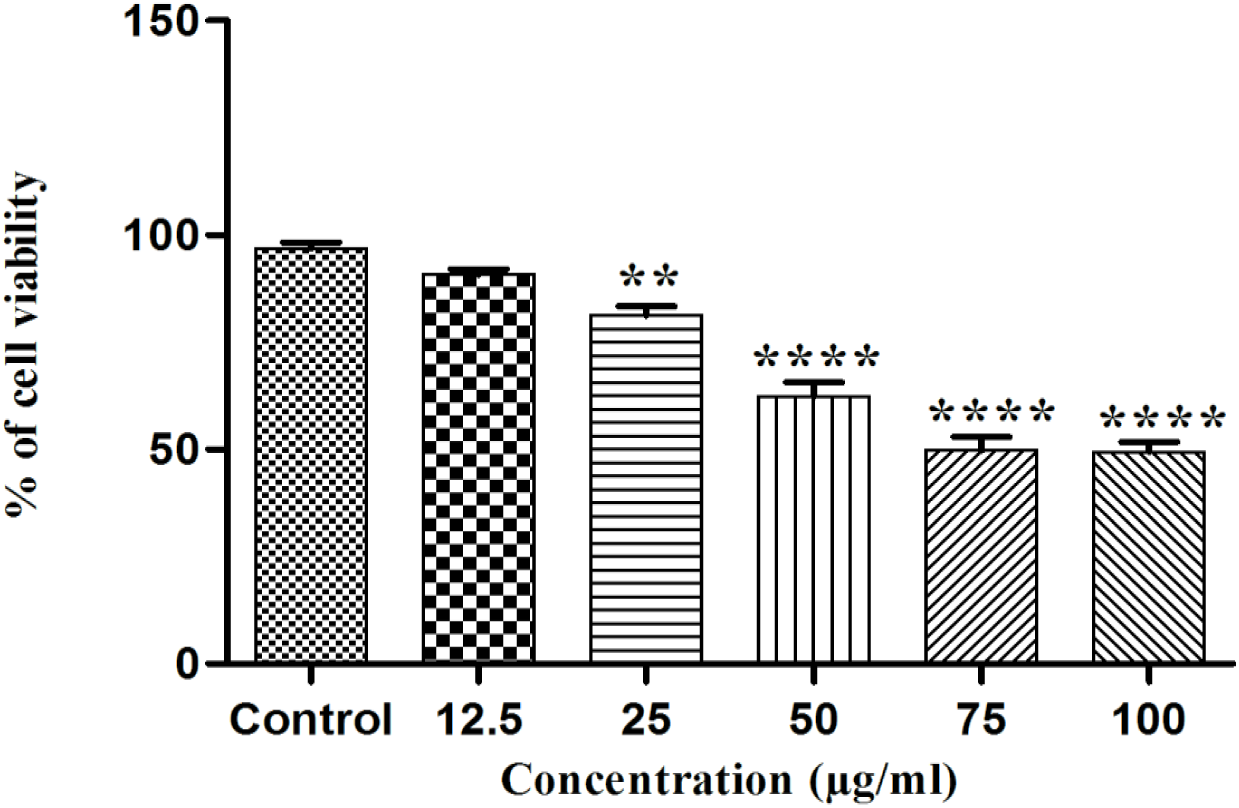
Concentration-dependent cell viability of methanol extract against PC_3_ cell line at 48 h. The PC_3_ cells were treated with leaf extracts (12.5, 25, 50, 75, 100 μg/mL) for 48 h. Results are presented as the mean ± SD of at least triplicates of each experiment. \**p* < 0.05, \*\**p* < 0.01, \*\*\**p* < 0.001 and \*\*\*\**p* < 0.0001, statistically significant in comparison with control groups by one-way ANOVA and followed by Bonferroni’s multiple comparison test

The dose dependent structural deformities in PC_3_ cell lines are represented in Figure 12 and Figure 14. Here, the result exhibited significant anti-proliferative effects of methanol extract. The remarkable cytotoxic effects of leaf extracts were characterized by the irreversible morphological changes of shapes of cells and dissociation from the surface of the plate at higher doses. However, these concentrations showed positive effects on cell viability of WI38 cell lines and recommended as favourable doses for assessing the cytotoxic activity on PC3 cell (Figure 9 and Figure 10).

**Figure 14:**
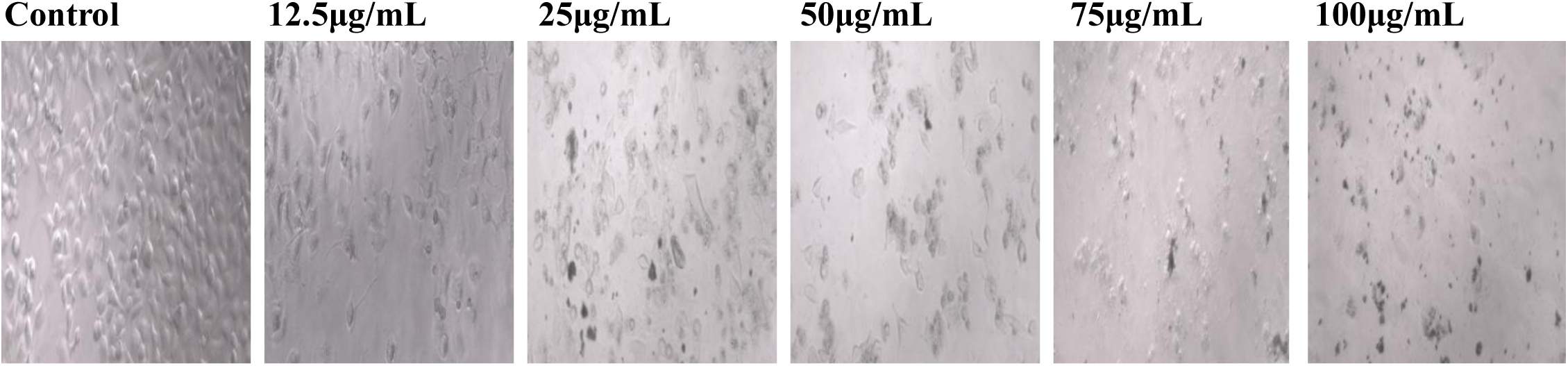
Concentration-dependent finite deformation effects in cell morphology of PC_3_. After treating with the 12.5 μg/mL, 25 μg/mL, 50 μg/mL, 75 μg/mL, 100μg/mL of methanol extract for 48 h

Numerous research reports support a significant correlation between polyphenols consumption and reduced risk of prostate cancer [65], [66]. Exposure to polyphenols may regulate the signalling cascades by increasing or decreasing some levels of proteins, producing pro-oxidant or antioxidant effects, stopping progression of cell cycle, and initiating the apoptosis pathway. For instance, apigenin has been found to be growth inhibitor of prostate cancer[67], [68] cells by pulling down the potential of mitochondrial membrane, and subsequently by producing the cytochrome c into cytoplasm, reducing the levels of anti-apoptotic proteins Bcl-2 and Bcl-2-extra-large (Bcl-XL) proteins and elevating the level of Bax. Additionally, apigenin plays a cardinal role in prevention of prostate cancer by involving TNF-related apoptosis-inducing ligand (TRAIL) and death receptor 5 (DR5) [64]. Ellagic acid is a well-known natural phenolic compound, and previous research have shown that ellagic acid is adequately present in our research sample [16]. Many of the research articles suggested the potent regulatory role of ellagic acid in prostate cancer [69], [70]. By impairing the activity of heme-oxygenase system, blocking the activity of vascular endothelial growth factor (VEGF) and osteoprotegerin (OPG), suppressing the expression of cyclin D1, and downregulating the levels of prostate-specific antigen (PSA), ellagic acid slows down the progression of metastasis and may induce the cell cycle arrest. Anthocyanin is a well-known subclass of flavonoids [71]. One of the major metabolites of anthocyanin is protocatechuic acid, and it is potentially employed for apoptotic cell death in prostate cancer [64]. Through enhanced caspase-3 activity, decreased VEGF activity, impaired the mitochondrial membrane potential and reduced the levels of pro-inflammatory cytokines (IL-6, IL-8), protocatechuic acid destroys the prostate cancer cells. In addition, it is well documented that, gallic acid exhibits chemoprevention against PC3 cells by lowering the levels of some proteins like, protein kinase C (PKC), NF-κB, c-Jun N-terminal kinase (JNK), ERK1/2, p38-MAPK, and p-Akt (Thr308, Ser 473), growth factor receptor bound protein 2 (GRB2), and by suppressing the gene expression of IL-6 [72].

### 3.7 Conclusion

In this research work, we explored the antioxidant potentials of the of *Vitex negundo* leaves collected in spring time from the West Bengal, India.

MAE procedure was used for robust extraction of phenolic and antioxidant compounds using less amount of solvent, energy and time. The BBD method worked well for maximizing the extraction of TPC, TFC and DPPH, ABTS from the leaves of *Vitex negundo* plant.

RSM was employed to get the optimum conditions (48 °C, 14 min, methanol proportion of 65% (v/v), and extraction volume of 14 mL) for MAE to extract the maximum amounts of polyphenols and antioxidants from the *Vitex negundo* leaves. These extraction settings yielded the TPC and TFC values of 1.464 ± 0.05 mg GAE/g extract and 1.065 ± 0.05 mg QE/g extract respectively. In addition, the antioxidant power was determined by the DPPH assay and ABTS assay and values were found to be 53.033 ± 0.32 and 77.720 ± 0.54 respectively.

The three-dimensional response surface plots vividly demonstrated that a little rise in methanol concentration and increased extraction volume, coupled with adequate extraction time and temperature, resulted in increased production of phenolic compounds and antioxidant capacity in the ideal extract.

GC-MS was employed to find out the 12 bioactive compounds from several classes including disaccharide, sesquiterpenoids, monoterpene, fatty acid ester, sesquiterpene, monocyclic diterpene alcohol, monoterpene aldehyde, diterpene alcohol, fatty acid, amino compound, triterpene; were inferred to be presented in the final extract. Furthermore, the presence of substantial amount of tocopherol proved that the *Vitex negundo* leaves is an excellent source of antioxidant. We also evaluated the anticancer effects of the leaf extract on PC3 cell lines. And the results depicted significant dose dependent cytotoxic effects on prostate carcinoma cell lines.

The present study comes up with proofs on the worth of *Vitex negundo* leaves as a great source natural antioxidant. The key role of obtained phyto-constituents including the antioxidant compounds is in alleviation from chronic disorder like cancer. A per the research findings, this medicinal plant can be a great therapeutic candidate in the near future, and could also be the subject of further scientific research.

## Footnotes

1 https://cran.r-project.org/web/packages/rsm/index.html

2 https://www.r-project.org/

## References

[1] S.-Y. Pan et al., ‘Historical Perspective of Traditional Indigenous Medical Practices: The Current Renaissance and Conservation of Herbal Resources’, Evidence-Based Complementary and Alternative Medicine, vol. 2014, pp. 1–20, 2014.

[2] N. Belkacem, B. Khettal, M. Hudaib, Y. Bustanji, B. Abu-Irmaileh, and C. S. M. Amrine, ‘Antioxidant, antibacterial, and cytotoxic activities of Cedrus atlantica organic extracts and essential oil’, Eur J Integr Med, vol. 42, p. 101292, Feb. 2021.

[3] F. Basri, H. P. Sharma, S. Firdaus, P. Jain, and A. Ranjan, ‘A review of ethnomedicinal plant-Vitex negundo Linn’, *Int*. J. Adv. Res, vol. 2, no. 3, pp. 882–894, 2014.

[4] A. Rani and A. Sharma, ‘The genus Vitex: A review’, Pharmacogn Rev, vol. 7, no. 14, p. 188, 2013.

[5] A. S. Vishwanathan and R. Basavaraju, ‘A review on Vitex negundo L.: A medicinally important plant’, Eur J Biol Sci, vol. 3, no. 1, pp. 30–42, 2010.

[6] G. Garg, A. Bharadwaj, S. Chaudhary, and V. Gupta, ‘Chemical profiling of bioactive compounds in the methanolic extract of wild leaf and callus of *Vitex negundo* using gas chromatography-mass spectrometry’, World J Exp Med, vol. 14, no. 1, Feb. 2024, doi: 10.5493/wjem.v14.i1.88064.

[7] V. Singh, R. Dayal, and J. Bartley, ‘Volatile Constituents of *Vitex negundo* Leaves’, Planta Med, vol. 65, no. 06, pp. 580–582, Feb. 1999, doi: 10.1055/s-2006-960832.

[8] Md. H. Al Rashid et al., ‘ANTIOXIDANT AND ANTICANCER ACTIVITY OF EXTRACT AND FRACTIONS OBTAINED FROM DIOSPYROS MELANOXYLON ROXB. LEAVES AND CORRELATION WITH THEIR POLYPHENOLIC PROFILES’, Int J Pharm Pharm Sci, vol. 10, no. 11, p. 6, Nov. 2018.

[9] O. P. Heinonen et al., ‘Prostate Cancer and Supplementation With α-Tocopherol and β-Carotene: Incidence and Mortality in a Controlled Trial’, JNCI: Journal of the National Cancer Institute, vol. 90, no. 6, pp. 440–446, Feb. 1998, doi: 10.1093/jnci/90.6.440.

[10] J. L. Watters, M. H. Gail, S. J. Weinstein, J. Virtamo, and D. Albanes, ‘Associations between α-Tocopherol, β-Carotene, and Retinol and Prostate Cancer Survival’, Cancer Res, vol. 69, no. 9, pp. 3833–3841, Feb. 2009, doi: 10.1158/0008-5472.CAN-08-4640.

[11] A. B. Awad, C. S. Fink, H. Williams, and U. Kim, ‘In vitro and in vivo (SCID mice) effects of phytosterols on the growth and dissemination of human prostate cancer PC-3 cells’, European Journal of Cancer Prevention, vol. 10, no. 6, pp. 507–513, Dec. 2001.

[12] B. Tohidi, M. Rahimmalek, and A. Arzani, ‘Essential oil composition, total phenolic, flavonoid contents, and antioxidant activity of Thymus species collected from different regions of Iran’, Food Chem, vol. 220, pp. 153–161, Apr. 2017.

[13] T. Rabi and S. Gupta, ‘Dietary terpenoids and prostate cancer chemoprevention’, Front Biosci, vol. 13, p. 3457, 2008.

[14] M. Z. M. Salem, H. M. Ali, and M. O. Basalah, ‘Essential oils from wood, bark, and needles of Pinus roxburghii Sarg. from Alexandria, Egypt: Antibacterial and antioxidant activities’, Bioresources, vol. 9, no. 4, pp. 7454–7466, 2014.

[15] M. Bin Sayeed, S. Karim, T. Sharmin, and M. Morshed, ‘Critical Analysis on Characterization, Systemic Effect, and Therapeutic Potential of Beta-Sitosterol: A Plant-Derived Orphan Phytosterol’, Medicines, vol. 3, no. 4, p. 29, Nov. 2016.

[16] A. Kundu, V. Mandal, P. P. Maiti, Md. H. Al Rashid, and S. C. Mandal, ‘In-vitro-Scientific evaluation of anti-inflammatory potential of leaf extracts from Vitex negundo: as a promising future drug candidate’, International Journal of Green Pharmacy (IJGP), vol. 14, no. 1, 2020.

[17] L. Abidin, M. Mujeeb, and S. R. Mir, ‘Maximized Extraction of Flavonoid Luteolin from V.negundo L. Leaves: Optimization Using Box-Behnken Design’, Curr Bioact Compd, vol. 15, no. 3, pp. 343–350, Feb. 2019, doi: 10.2174/1573407214666180731120014.

[18] J. Cheng et al., ‘pH-responsive functionalized surface active ionic liquid as an enhanced medium for efficient extraction and in-situ separation of flavonoids in Vitex negundo L. Leaves’, Microchemical Journal, vol. 193, p. 109080, Feb. 2023, doi: 10.1016/j.microc.2023.109080.

[19] A. kumar Katare et al., ‘Optimisation of Extraction Process for Negundoside and Agnuside from Vitex Negundo L. Leaves Using Soxhlet Extraction, HPLC–MS/MS, and CCD-RSM Methods’, CHEMISTRY AFRICA-A JOURNAL OF THE TUNISIAN CHEMICAL SOCIETY, vol. 5, no. 4, pp. 907–915, Feb. 2022.

[20] V. Ummat et al., ‘Optimisation of Ultrasound Frequency, Extraction Time and Solvent for the Recovery of Polyphenols, Phlorotannins and Associated Antioxidant Activity from Brown Seaweeds’, Mar Drugs, vol. 18, no. 5, p. 250, Feb. 2020, doi: 10.3390/md18050250.

[21] N. A. Latiff, P. Y. Ong, S. N. A. A. Rashid, L. C. Abdullah, N. A. M. Amin, and N. A. M. Fauzi, ‘Enhancing recovery of bioactive compounds from Cosmos caudatus leaves via ultrasonic extraction’, Sci Rep, vol. 11, no. 1, p. 17297, Feb. 2021, doi: 10.1038/s41598-021-96623-x.

[22] A. G. Demesa, S. Saavala, M. Pöysä, and T. Koiranen, ‘Overview and Toxicity Assessment of Ultrasound-Assisted Extraction of Natural Ingredients from Plants’, Foods, vol. 13, no. 19, p. 3066, Feb. 2024, doi: 10.3390/foods13193066.

[23] A. S. L. Pérez, J. J. L. Castro, and C. A. G. Fajardo, ‘Application of Microwave Energy to Biomass: A Comprehensive Review of Microwave-Assisted Technologies, Optimization Parameters, and the Strengths and Weaknesses’, Journal of Manufacturing and Materials Processing, vol. 8, no. 3, p. 121, Feb. 2024, doi: 10.3390/jmmp8030121.

[24] B. Kaufmann and P. Christen, ‘Recent extraction techniques for natural products: microwave-assisted extraction and pressurised solvent extraction’, Phytochemical Analysis, vol. 13, no. 2, pp. 105–113, Feb. 2002, doi: 10.1002/pca.631.

[25] A. Khalfi, M. C. Garrigós, M. Ramos, and A. Jiménez, ‘Optimization of the Microwave-Assisted Extraction Conditions for Phenolic Compounds from Date Seeds’, Foods, vol. 13, no. 23, p. 3771, Feb. 2024, doi: 10.3390/foods13233771.

[26] Z. Karami, Z. Emam-Djomeh, H. A. Mirzaee, M. Khomeiri, A. S. Mahoonak, and E. Aydani, ‘Optimization of microwave assisted extraction (MAE) and soxhlet extraction of phenolic compound from licorice root’, J Food Sci Technol, Feb. 2014, doi: 10.1007/s13197-014-1384-9.

[27] K. Ameer, S.-W. Bae, Y. Jo, H.-G. Lee, A. Ameer, and J.-H. Kwon, ‘Optimization of microwave-assisted extraction of total extract, stevioside and rebaudioside-A from Stevia rebaudiana (Bertoni) leaves, using response surface methodology (RSM) and artificial neural network (ANN) modelling’, Food Chem, vol. 229, pp. 198–207, Feb. 2017, doi: 10.1016/j.foodchem.2017.01.121.

[28] G. Aquino et al., ‘Optimization of microwave-assisted extraction of antioxidant compounds from spring onion leaves using Box–Behnken design’, Sci Rep, vol. 13, no. 1, p. 14923, Feb. 2023, doi: 10.1038/s41598-023-42303-x.

[29] Y. Ma, J. Li, F. Tong, X.-L. Xin, and H. A. Aisa, ‘Optimization of microwave-assisted extraction using response surface methodology and the potential anti-diabetic efficacy of Nigella glandulifera Freyn determined using the spectrum–effect relationship’, Ind Crops Prod, vol. 153, p. 112592, Feb. 2020, doi: 10.1016/j.indcrop.2020.112592.

[30] H. K. Kala, R. Mehta, K. K. Sen, R. Tandey, and V. Mandal, ‘Strategizing method optimization of microwave-assisted extraction of plant phenolics by developing standard working principles for universal robust optimization’, Analytical Methods, vol. 9, no. 13, pp. 2089–2103, 2017.

[31] K. B. S. Chouhan, R. Tandey, K. K. Sen, R. Mehta, and V. Mandal, ‘Extraction of phenolic principles: Value addition through effective sample pretreatment and operational improvement’, Journal of Food Measurement and Characterization, vol. 13, no. 1, pp. 177–186, 2019.

[32] M. Ondua, E. M. Njoya, M. A. Abdalla, and L. J. McGaw, ‘Anti-inflammatory and antioxidant properties of leaf extracts of eleven South African medicinal plants used traditionally to treat inflammation’, J Ethnopharmacol, vol. 234, pp. 27–35, Apr. 2019.

[33] Y. Ge, H. Yan, B. Hui, Y. Ni, S. Wang, and T. Cai, ‘Extraction of Natural Vitamin E from Wheat Germ by Supercritical Carbon Dioxide’, J Agric Food Chem, vol. 50, no. 4, pp. 685–689, Feb. 2002.

[34] V. K. Nelson et al., ‘Azadiradione ameliorates polyglutamine expansion disease in Drosophila by potentiating DNA binding activity of heat shock factor 1’, Oncotarget, vol. 7, no. 48, p. 78281, 2016.

[35] N. F. Azahar, S. S. A. Gani, and N. F. M. Mokhtar, ‘Optimization of phenolics and flavonoids extraction conditions of Curcuma Zedoaria leaves using response surface methodology’, Chem Cent J, vol. 11, no. 1, p. 96, Feb. 2017, doi: 10.1186/s13065-017-0324-y.

[36] F. Imtiaz, D. Ahmed, R. H. Abdullah, and S. Ihsan, ‘Green extraction of bioactive compounds from Thuja orientalis leaves using microwave- and ultrasound-assisted extraction and optimization by response surface methodology’, Sustain Chem Pharm, vol. 35, p. 101212, Feb. 2023, doi: 10.1016/j.scp.2023.101212.

[37] A. Mehmood, S. Javid, M. F. Khan, K. S. Ahmad, and A. Mustafa, ‘In vitro total phenolics, total flavonoids, antioxidant and antibacterial activities of selected medicinal plants using different solvent systems’, BMC Chem, vol. 16, no. 1, p. 64, Feb. 2022, doi: 10.1186/s13065-022-00858-2.

[38] A. Wakeel, S. A. Jan, I. Ullah, Z. K. Shinwari, and M. Xu, ‘Solvent polarity mediates phytochemical yield and antioxidant capacity of *Isatis tinctoria*’, PeerJ, vol. 7, p. e7857, Feb. 2019, doi: 10.7717/peerj.7857.

[39] A. H. Nour, A. R. Oluwaseun, A. H. Nour, M. S. Omer, and N. Ahmed, ‘Microwave-Assisted Extraction of Bioactive Compounds (Review)’, in *Microwave Heating - Electromagnetic Fields Causing Thermal and Non-Thermal Effects*, IntechOpen, 2021. doi: 10.5772/intechopen.96092.

[40] J. F. Osorio-Tobón, ‘Recent advances and comparisons of conventional and alternative extraction techniques of phenolic compounds’, J Food Sci Technol, vol. 57, no. 12, pp. 4299–4315, Feb. 2020, doi: 10.1007/s13197-020-04433-2.

[41] A. Antony and M. Farid, ‘Effect of Temperatures on Polyphenols during Extraction’, Applied Sciences, vol. 12, no. 4, p. 2107, Feb. 2022, doi: 10.3390/app12042107.

[42] X. Liu et al., ‘Effects of five extraction methods on total content, composition, and stability of flavonoids in jujube’, Food Chem X, vol. 14, p. 100287, Feb. 2022, doi: 10.1016/j.fochx.2022.100287.

[43] Y. Gupta, B. Barrett, and D. G. Vlachos, ‘Understanding microwave-assisted extraction of phenolic compounds from diverse food waste feedstocks’, Chemical Engineering and Processing - Process Intensification, vol. 203, p. 109870, Feb. 2024, doi: 10.1016/j.cep.2024.109870.

[44] R. Albarri, İ. Toprakçı, E. Kurtulbaş, and S. Şahin, ‘Estimation of diffusion and mass transfer coefficients for the microwave-assisted extraction of bioactive substances from Moringa oleifera leaves’, Biomass Convers Biorefin, vol. 13, no. 6, pp. 5125–5132, Feb. 2023, doi: 10.1007/s13399-021-01443-8.

[45] W. Elfalleh, B. Kirkan, and C. Sarikurkcu, ‘Antioxidant potential and phenolic composition of extracts from Stachys tmolea: An endemic plant from Turkey’, Ind Crops Prod, vol. 127, pp. 212–216, Jan. 2019.

[46] D. Tungmunnithum, A. Thongboonyou, A. Pholboon, and A. Yangsabai, ‘Flavonoids and Other Phenolic Compounds from Medicinal Plants for Pharmaceutical and Medical Aspects: An Overview’, Medicines, vol. 5, no. 3, p. 93, Aug. 2018.

[47] Z. Réblová, ‘Effect of temperature on the antioxidant activity of phenolic acids’, Czech Journal of Food Sciences, vol. 30, no. 2, pp. 171–175, Feb. 2012, doi: 10.17221/57/2011-CJFS.

[48] E. Gil-Martín, T. Forbes-Hernández, A. Romero, D. Cianciosi, F. Giampieri, and M. Battino, ‘Influence of the extraction method on the recovery of bioactive phenolic compounds from food industry by-products’, Food Chem, vol. 378, p. 131918, Feb. 2022, doi: 10.1016/j.foodchem.2021.131918.

[49] R. R. Kotha, F. S. Tareq, E. Yildiz, and D. L. Luthria, ‘Oxidative Stress and Antioxidants—A Critical Review on In Vitro Antioxidant Assays’, Antioxidants, vol. 11, no. 12, p. 2388, 2022, doi: 10.3390/antiox11122388.

[50] V. Mareček et al., ‘ABTS and DPPH methods as a tool for studying antioxidant capacity of spring barley and malt’, J Cereal Sci, vol. 73, pp. 40–45, Jan. 2017.

[51] D. Mukhopadhyay et al., ‘A Sensitive In vitro Spectrophotometric Hydrogen Peroxide Scavenging Assay using 1,10-Phenanthroline’, Free Radicals and Antioxidants, vol. 6, no. 1, pp. 124–132, Jan. 2016.

[52] G. Derringer and R. Suich, ‘Simultaneous Optimization of Several Response Variables’, Journal of Quality Technology, vol. 12, no. 4, pp. 214–219, Feb. 1980, doi: 10.1080/00224065.1980.11980968.

[53] J. Legault and A. Pichette, ‘Potentiating effect of β-caryophyllene on anticancer activity of α-humulene, isocaryophyllene and paclitaxel’, Journal of Pharmacy and Pharmacology, vol. 59, no. 12, pp. 1643–1647, Feb. 2010.

[54] E. B. Russo, ‘Taming THC: potential cannabis synergy and phytocannabinoid-terpenoid entourage effects’, Br J Pharmacol, vol. 163, no. 7, pp. 1344–1364, Aug. 2011.

[55] H. B. Martins et al., ‘Anti-Inflammatory Activity of the Essential Oil Citral in Experimental Infection with Staphylococcus aureus in a Model Air Pouch’, Evidence-Based Complementary and Alternative Medicine, vol. 2017, pp. 1–10, 2017.

[56] H. Sales-Campos, P. Reis de Souza, B. Crema Peghini, J. Santana da Silva, and C. Ribeiro Cardoso, ‘An Overview of the Modulatory Effects of Oleic Acid in Health and Disease’, Mini Rev Med Chem, vol. 13, no. 2, pp. 201–210, Feb. 2013.

[57] S. Muthukumarasamy, V. R. Mohan, and others, ‘GC-MS determination of bioactive components of Canscora perfoliata Lam.(Gentianaceae)’, J Appl Pharm Sci, vol. 2, no. 8, pp. 210–214, 2012.

[58] R. Chelliah, S. Ramakrishnan, and U. Antony, ‘Nutritional quality of Moringa oleifera for its bioactivity and antibacterial properties’, Int Food Res J, vol. 24, no. 2, p. 825, 2017.

[59] G. Sethi, S. Sankaranarayanan, and M. Sukhija, ‘Low lactose in the nutritional management of diarrhea: Case reports from India’, Clin Epidemiol Glob Health, vol. 6, no. 4, pp. 160–162, Dec. 2018.

[60] P. Arulselvan et al., ‘Role of Antioxidants and Natural Products in Inflammation’, Oxid Med Cell Longev, vol. 2016, pp. 1–15, 2016.

[61] F. Al-Rimawi et al., ‘Anticancer, antioxidant, and antibacterial activity of chemically fingerprinted extract from Cyclamen persicum Mill.’, Sci Rep, vol. 14, no. 1, p. 8488, Feb. 2024, doi: 10.1038/s41598-024-58332-z.

[62] S. Jongrungraungchok et al., ‘In vitro antioxidant, anti-inflammatory, and anticancer activities of mixture Thai medicinal plants’, BMC Complement Med Ther, vol. 23, no. 1, p. 43, Feb. 2023, doi: 10.1186/s12906-023-03862-8.

[63] S. Sun, Z. Liu, M. Lin, N. Gao, and X. Wang, ‘Polyphenols in health and food processing: antibacterial, anti-inflammatory, and antioxidant insights’, Front Nutr, vol. 11, Feb. 2024, doi: 10.3389/fnut.2024.1456730.

[64] B. Gullón, G. Eibes, M. T. Moreira, I. Dávila, J. Labidi, and P. Gullón, ‘Antioxidant and antimicrobial activities of extracts obtained from the refining of autohydrolysis liquors of vine shoots’, Ind Crops Prod, vol. 107, pp. 105–113, Nov. 2017.

[65] Y. Huang, W. Wang, and J. Jin, ‘Association between polyphenol subclasses and prostate cancer: a systematic review and meta-analysis of observational studies’, Front Nutr, vol. 11, 2024, doi: 10.3389/fnut.2024.1428911.

[66] T. Costea, P. Nagy, C. Ganea, J. Szöllősi, and M.-M. Mocanu, ‘Molecular Mechanisms and Bioavailability of Polyphenols in Prostate Cancer’, Int J Mol Sci, vol. 20, no. 5, p. 1062, Feb. 2019, [Online]. Available: https://www.mdpi.com/1422-0067/20/5/1062

[67] A. H. Rahmani et al., ‘The Potential Role of Apigenin in Cancer Prevention and Treatment’, Molecules, vol. 27, no. 18, p. 6051, 2022, doi: 10.3390/molecules27186051.

[68] S. Shukla, N. Bhaskaran, M. A. Babcook, P. Fu, G. T. MacLennan, and S. Gupta, ‘Apigenin inhibits prostate cancer progression in TRAMP mice via targeting PI3K/Akt/FoxO pathway’, Carcinogenesis, vol. 35, no. 2, pp. 452–460, 2014, doi: 10.1093/carcin/bgt316.

[69] P. Pitchakarn et al., ‘Ellagic Acid Inhibits Migration and Invasion by Prostate Cancer Cell Lines’, Asian Pacific Journal of Cancer Prevention, vol. 14, no. 5, pp. 2859–2863, 2013, doi: 10.7314/APJCP.2013.14.5.2859.

[70] A. Naiki-Ito et al., ‘Ellagic acid, a component of pomegranate fruit juice, suppresses androgen-dependent prostate carcinogenesis via induction of apoptosis’, Prostate, vol. 75, no. 2, pp. 151–160, Feb. 2015, [Online]. Available: https://onlinelibrary.wiley.com/doi/10.1002/pros.22900

[71] X. Zhou et al., ‘Analysis of anthocyanins and total flavonoids content in functional rice and its recombination inbred lines’, Front Plant Sci, vol. 14, Feb. 2023, doi: 10.3389/fpls.2023.1113618.

[72] E. Heidarian, M. Keloushadi, K. Ghatreh-Samani, and P. Valipour, ‘The reduction of IL-6 gene expression, pAKT, pERK1/2, pSTAT3 signaling pathways and invasion activity by gallic acid in prostate cancer PC3 cells’, Biomedicine & Pharmacotherapy, vol. 84, pp. 264–269, Feb. 2016, doi: 10.1016/j.biopha.2016.09.046.

